# Addressing Erroneous Scale Assumptions in Microbe and Gene Set Enrichment Analysis

**DOI:** 10.1101/2023.03.10.532120

**Authors:** Kyle C. McGovern, Michelle Pistner Nixon, Justin D. Silverman

## Abstract

By applying Differential Set Analysis (DSA) to sequence count data, researchers can determine whether groups of microbes or genes are differentially enriched. Yet these data lack information about the scale (i.e., size) of the biological system under study, leading some authors to call these data compositional (i.e., proportional). In this article we show that commonly used DSA methods make strong, implicit assumptions about the unmeasured system scale. We show that even small errors in these assumptions can lead to false positive rates as high as 70%. To mitigate this problem, we introduce a sensitivity analysis framework to identify when modeling results are robust to such errors and when they are suspect. Unlike standard benchmarking studies, our methods do not require ground-truth knowledge and can therefore be applied to both simulated and real data.

## 1 Introduction

Sequence count data (e.g., data from 16S rRNA-seq or RNA-seq studies) have become ubiquitous in modern biomedical research. These data are often used to identify whether a pre-determined set of entities (i.e., set of microbes or set of genes) are differentially enriched between two biological conditions [1, 2]. For example, in the analysis of RNA-Seq data, researchers often use tools such as Gene Set Enrichment Analysis (GSEA) [1] to identify pathways (sets of genes) that are up or down regulated between conditions [3–5]. We refer to this inferential task as Differential Set Analysis (DSA). Here we show that common approaches to DSA can display high rates of false positives due to limitations of the sequence counting measurement process. We introduce new tools and insights to help researchers mitigate these errors.

Sequencing depth (the number of measured DNA molecules in a sample) can vary substantially between samples due to non-biological (i.e., technical) factors [6, 7]. As a result, many authors use some form of statistical normalization, with the goal of removing unwanted variation and facilitating inference that requires between sample comparisons [8, 9]. Others argue for a different perspective: rather than viewing this non-biological variation as *extra noise* that must be removed, they view it as a symptom of an imperfect measurement process that *lacks information* about the scale (i.e., total size) of the underlying biological system [7, 10, 11]. Following this logic, the information contained in the data appears fundamentally compositional (relative), leading some authors to use the principle of scale invariance from Compositional Data Analysis (CoDA) to guide analyses [7, 10]. The scale invariance principle restricts analyses to those that are invariant to the system scale [12]: analyses for which the lack of scale information is immaterial. While intuitive, this approach can be scientifically limiting, as many biologically relevant scientific questions require scale information [13, 14].

A third perspective has recently emerged. Nixon et al. proposed a reformulation of CoDA called *Scale Reliant Inference* (SRI) for statistically rigorous non-scale invariant (scale reliant) analyses with sequence count data [11]. Using SRI, they showed that many methods for analyzing sequence count data attempt to overcome the lack of scale information by using strong, implicit assumptions about the unmeasured system scale – scale assumptions [11]. By applying their framework to the study of Differential Analysis (DA, i.e., differential abundance and differential expression analyses) they show that even slight errors in scale assumptions can lead to false discovery rates as high as 80%. Viewing DSA as a generalization of DA, where the goal is to study sets of entities rather than individual entities, we hypothesized that errors in scale assumptions may lead to inferential errors in DSA.

We review concepts and terminology from the field of SRI to study how errors in scale assumptions can impact DSA. As in SRI, our approach centers around a mathematical quantity called a *target estimand*, the quantity a researcher wants to estimate. We study the impact of erroneous scale assumptions and characterize how this impact depends on the research goal. We show that many common research goals are scale reliant and that, in the context of those goals, common methods for performing DSA result in elevated rates of false positives. To mitigate these problems, we develop a type of sensitivity analysis to identify possible false positives. In addition, we characterize a broad class of research goals (target estimands) that are insensitive to erroneous scale assumptions, and discuss how to determine if research goals fall within this class. Beyond providing new tools for performing DSA, the overall purpose of this article is to catalyze future research and discussions into the impact of scale limitations in sequence count data; to show that rigorous and reproducible scale reliant inference is possible with these data; and to also highlight that care is required to perform such analyses.

### 1.1 Review of Scale Reliant Inference

Scale Reliant Inference (SRI) is a conceptual framework to study scale assumptions in sequence count data [11]. We review key terms and concepts from SRI relevant to our study of DSA. For concreteness, we present this material in terms of a hypothetical study performing differential abundance analysis on a cross-sectional microbiome dataset of *N* study participants, half with a disease of interest and half healthy controls.

#### 1.1.1 The Scaled System, Observed Data, and Target Estimand

For differential abundance we represent the observed data as a *D × N* matrix of counts *Y* with elements *Y*_*dn*_ representing the number of DNA molecules that map to the *d*-th microbial taxon in the sample obtained from participant *n*. Central to SRI is the notion that the observed data (*Y*) is a measurement of an underlying biological system called a *scaled system* and denoted as *W*. Like *Y, W* is a *D × N* matrix. Unlike *Y*, the elements *W*_*dn*_ represent the true (as opposed to measured) amount of the *d*-th taxon in the gut microbiota of the *n*-th participant. Importantly, note that the definition of *true amount* is not restricted to the total number of microbes in a subject’s gut. For example, for a researcher interested in host-pathogen interactions, true amounts could be the ratio of bacterial to human cells at the epithelial surface of the distal colon.

The scaled system represents an unmeasured standard of truth, a reference against which we can discuss the limitations of the observed data. While the sequence counting process can provide limited information about the scaled system in many ways (e.g., PCR bias [15]), this article studies the lack of scale information in these data. Consider that the scaled system can be uniquely described in terms of its scale (summed, *W*^⊥^) and compositional (proportional, *W*^∥^) parts:

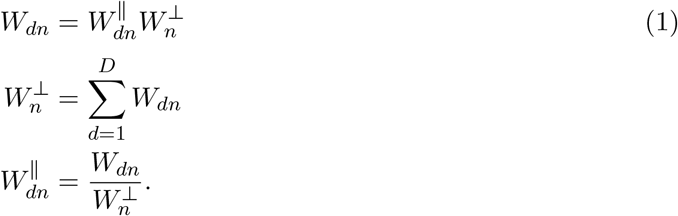

Intuitively, saying sequence count data lacks scale information means that, due to the non-biological variation in sequencing depth, sequence count data cannot be used to estimate the system scale (*W*^⊥^) to any reasonable degree of precision [8, 9] (see Nixon et al. for a more formal definition involving model identifiability [11]).

Ultimately, the extent to which scale limitations matter to an applied researcher depends on their research goal. In SRI, the research goal is represented by a mathematical quantity called a target estimand. Formally, a target estimand (*ψ*) is the output of a target estimator (a function *f*) applied to the scaled system: *ψ* = *f* (*W*). For example, a differential abundance analysis may look at the Log-Fold-Change (LFC) of each taxa *d*:

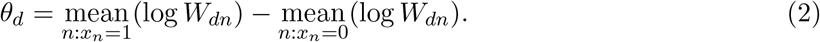

for a binary covariate *x*_*n*_ denoting if a participant is healthy or diseased.

The fundamental challenge of SRI occurs when the desired target estimand relies upon scale information not present in the observed data. For example, using the relationship log 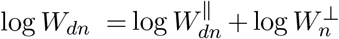 implied by equation (1), the LFC target estimand in equation (2) can be written as

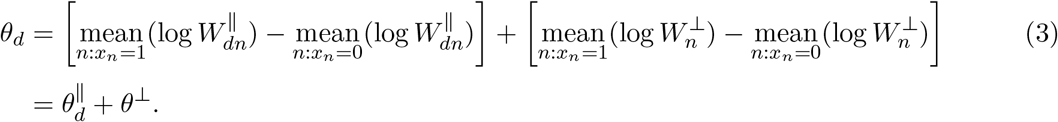

This target estimand is scale reliant: it requires knowledge of *θ*^⊥^ and therefore *W*^⊥^ to uniquely determine *θ*_*d*_ (see Nixon, et al. for a formal definition in terms of the invertibility of maps [11]).

#### 1.1.2 Scale Assumptions and the Applied Estimator

The target estimand and estimator, the scaled system, and a compositional survey make up the three core constructs of SRI. From this core, we explicitly define the goal of an analysis and the challenge of achieving that goal given the limited information content of the observed data. Notably, this core does not include the specific analytical method applied. That method, referred to as an applied estimator *g*, is a function applied to the observed data to estimate the target estimand 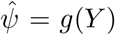. In SRI, the target estimand and estimator serve as a backdrop against which we can study the assumptions and quantify errors made by the applied estimator. For example, consider observed data (*Y*) that lacks scale information and a target estimand (*ψ*) that is scale reliant (that requires scale information). Against this backdrop, it is clear that *any* applied estimator 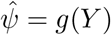 that produces a unique estimate of 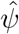 must have made some assumption about the unmeasured scale of the system *W*^⊥^. We call such assumptions *scale assumptions*.

Following Nixon et al. [11], we illustrate the concept of scale assumptions by studying CLR normalization in differential abundance analysis. Recognizing the scale limitations of sequence count data, a variety of tools (e.g., ALDEx2 [16] or Songbird [17]) estimate LFCs using Center Log-Ratio (CLR) normalized abundances. In brief, these methods can be thought of as producing estimates of a target estimand:

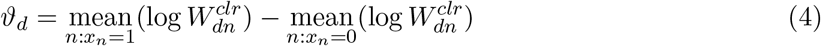

where 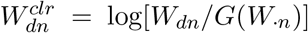 and where *G*(*W*_·*n*_) denotes the geometric mean of the vector *W*_·*n*_ = (*W*_1*n*_, …, *W*_*Dn*_). The target estimand in equation 4 is not the same as the LFC target estimand given in equation 2. But in our experience, many people take the estimates produced by ALDEx2 and Songbird as estimates of LFCs as defined in equation 2. Equating these two target estimators, i.e., assuming that the output of ALDEx2 or Songbird are estimates of LFCs as defined in equation (2) can be represented as the assumption that *θ*_*d*_ = *ϑ*_*d*_. We call this the CLR Assumption. We simplify this assumption to highlight the implied fixed relationship between the system composition and the system scale:

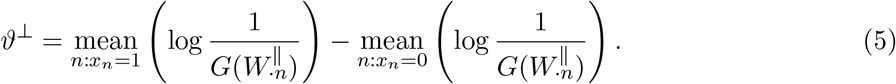

## 2 Results

### 2.1 Applied and Target Estimators for DSA

DSA infers, for a set of entities *S*, a quantity *ϕ*_*S*_ which can take on three values: +1 if the set is differentially enriched, −1 if differentially depleted, and 0 if neither enriched nor depleted. Applied estimators produce estimates of *ϕ*_*S*_ using observed data *Y*, while target estimators precisely delineate differential enrichment or depletion.

There are many applied estimators for DSA. A particularly popular approach is to apply Gene Set Enrichment Analysis (GSEA) [1] to LFCs estimated using tools such as ALDEx2 [16], Songbird [17], or DESeq2 [18]. Broadly, we call this general approach the GSEA-LFC applied estimator. At its core, GSEA-LFC uses GSEA to test whether the LFCs of entities in the set cluster together non-randomly when compared to the LFCs of entities not in the set. DSA estimators such as GSEALFC and others [19] can be thought of as two-stage applied estimators: the first stage estimator (e.g., ALDEx2,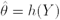) is responsible for estimating LFCs, and the second stage estimator uses those estimates to then estimate 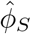(e.g., GSEA, 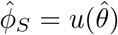). We next use this two-stage form to define target estimands that reflect common research goals in the field.

It is challenging to identify target estimators and estimands in practice. Just as researchers’ goals can differ, so too can their definitions of enriched and depleted entity sets; existing literature often does not explicitly state the estimation goals of a study. To address this challenge, we assume a correspondence between the applied estimator a researcher uses and their research goals. In the following sections, consider a researcher who uses a DSA applied estimator with second stage 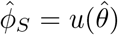. We assume the target estimand and estimator are given by 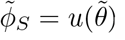, where we use a 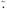 to explicitly denote the true value of a quantity.

### 2.2 LFC Sensitivity Analysis

In this section we describe a simple and computationally efficient sensitivity analysis framework to study how errors in scale assumptions impact DSA. We focus on DSA applied estimators and corresponding target estimators with the two stage form described above. For brevity, we generalize this approach to estimators that do not have this form (e.g., CAMERA [20]) in Supplementary Section 1.

Consider an applied estimator of the form 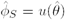 with corresponding target estimator and target estimand 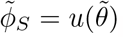. For any true value of the target estimand 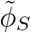 and any estimate 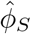 we represent the relationship between the two as 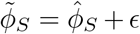 where *ϵ* represents error in the estimate. Following equation (3), we denote the LFC estimate as *θ* = *θ*^∥^ + *θ*^⊥^. Similarly, we think of the error *ϵ* as having compositional and scale parts: *ϵ* = *ϵ*^∥^ + *ϵ*^⊥^ where 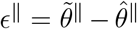 denotes error in the compositional component of the estimate and 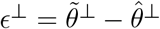 denotes error in the scale component. Since our goal is to understand the sensitivity of estimates to errors in scale assumptions, and since errors in scale assumptions only enter through the term *ϵ*^⊥^, we assume that *ϵ*^∥^ ≪ *ϵ*^⊥^ such that *ϵ* ≈ *ϵ*^⊥^. Then the true value 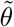 can be expressed in terms of the estimate 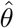 as

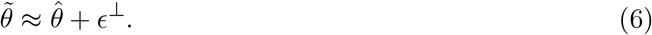

Returning to DSA, equation (6) can then be used to represent the truth 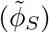 in terms of the estimates 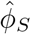:

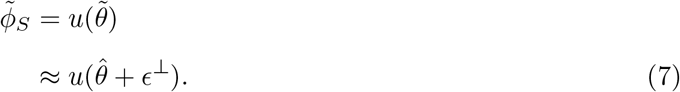

As we already know 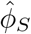, the output of the applied estimator, with equation (7) we can see how our estimates diverge from the truth 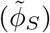 as a function of errors in scale assumptions (*ϵ*^⊥^). We call this LFC Sensitivity Analysis.

LFC Sensitivity Analysis has a number of appealing properties. To perform LFC sensitivity analysis, we only need to know the applied LFC estimates 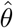 and not the true value of the DSA target estimand 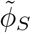 or the true value of the LFCs 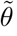. Thus this method can be applied to both simulated and real data.

Another appealing quality of LFC Sensitivity Analysis is its interpretability. Considering that 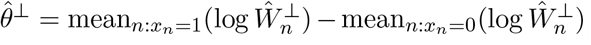, it follows that *ϵ*^⊥^ can be interpreted as the error in the log-fold-change of scales. For example, an error of *ϵ*^⊥^ = 1 can be read as a statement: *the ratio of the mean scale in the case condition compared to the control condition is e*^1^ ≈ 2.7 *times higher than assumed*.^1^ Moreover, this interpretation does not depend on the chosen notion of *amount* : any units attached to a researchers chosen notion of amount (e.g., cells per mL) cancel in the ratio.

### 2.3 The GSEA-LFC Target Estimand is Typically Scale Reliant

We chose to study GSEA-LFC target and applied estimators due to their popularity. The GSEA algorithm is visualized in Figure 1 and a formal definition is provided in Supplementary Section 2. In brief, GSEA is an algorithm that first orders entities by a ranking statistic (here LFCs) and then performs a hypothesis test to determine if entities in the set of interest cluster non-randomly at the top or bottom of that ordered list. The output of GSEA can be summarized as the quantity *ϕ*_*S*_ introduced above. Two key variations on the GSEA algorithm differ in the null model used to test the significance of the estimated 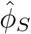 statistic. The first, which we highlight in the main text, permutes the entity labels. The second permutes the sample labels. Assuming adequate sample size, sample-label permutations are generally preferred, as they ensure that the null model accounts for inter-entity correlations [20–22]. However, sensitivity analyses when using sample-label permutations can be more complex; we leave them to Supplementary Section 1. That section also demonstrates that core conclusions of this work hold beyond GSEA and still apply to other popular DSA methods such as CAMERA [20].

**Figure 1:**
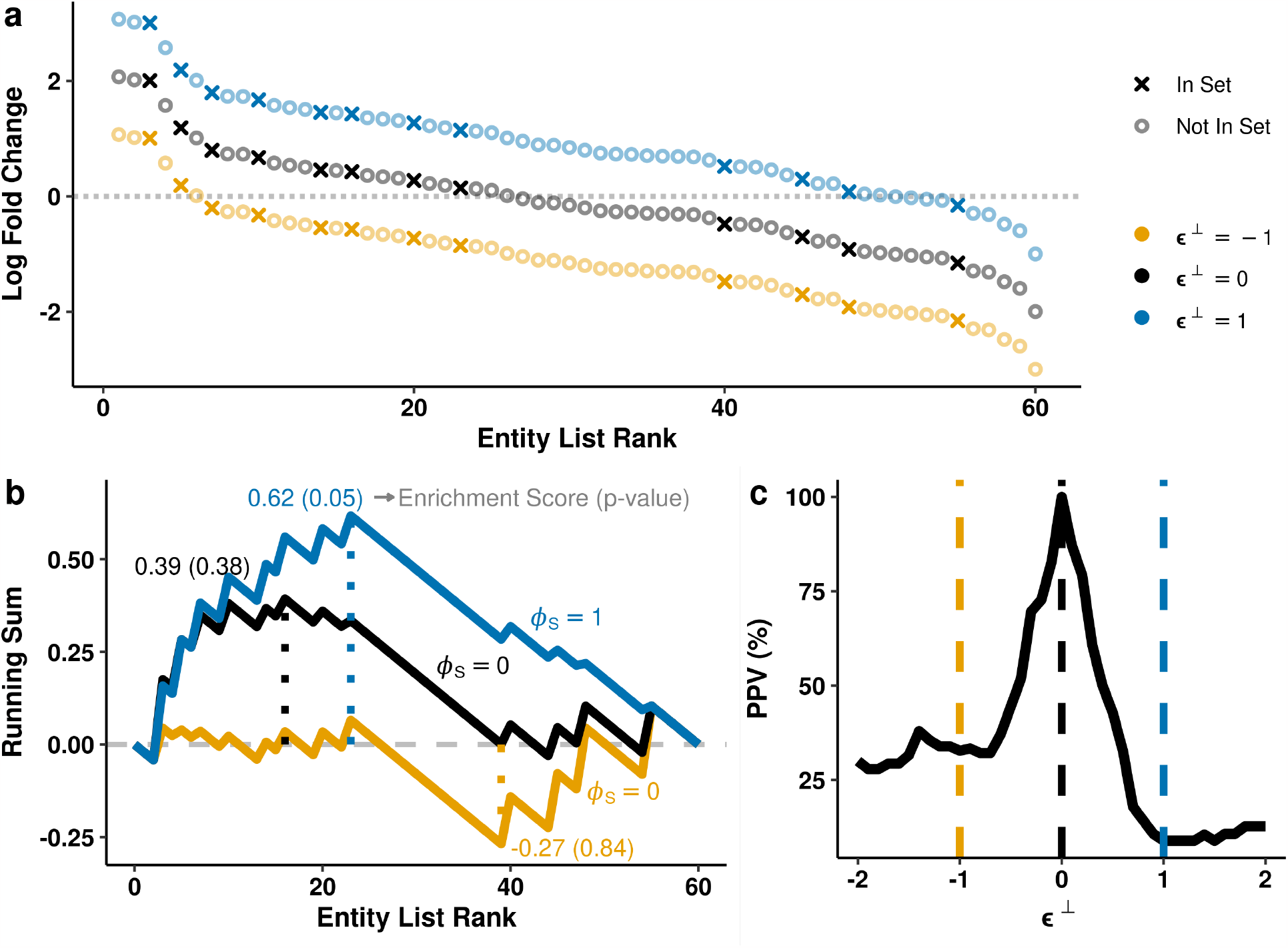
The GSEA-LFC target estimand is scale reliant. **(a)** The LFCs of 60 entities were simulated from a standard normal distribution and rank ordered (black); 12 of these entities were randomly selected to be in the entity set. For this simulation, the CLR assumption equated to a value of 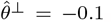. Blue and orange points represent the true LFCs if the assumed value of 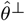 had error 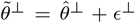. **(b)** For each level of error *ϵ*^⊥^ depicted in Panel a, we show the GSEA running sum and the corresponding enrichment scores (ES). The dotted lines represents the GSEA enrichment score, which is the maximum distance from zero of the running sum. P-values and *ϕ*_*S*_ values are also shown, although they depend on permutation tests which are not depicted visually. *ϕ*_*S*_ is the GSEA-LFC estimand. *ϕ*_*S*_ is ±1 when the entity set is significantly enriched or depleted (blue line, here *α* ≤ 0.05 is used for significance) and 0 when the entity set is not enriched (black and orange lines). **(c)** 10,000 entity sets of size 12 were simulated using the procedure as in Panel a in order to demonstrates how the Positive Predictive Value (PPV) of GSEA-LFC can vary with *ϵ*^⊥^. The dashed lines indicate the same values of *ϵ*^⊥^ shown in Panels a and b.

We used LFC Sensitivity Analysis to demonstrate that, for many entity sets, the GSEA-LFC target estimand is scale reliant. An example is depicted visually in Figure 1 which shows that different errors *ϵ*^⊥^ in the assumed scale 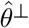 lead to different values of the target estimand 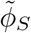. That is, for the depicted entity set, knowledge of the system composition alone is insufficient to uniquely determine the value of *ϕ*_*S*_.

To confirm that these results were not unique to the simulated entity set shown in Figure 1a and b, we repeated this analysis with 10,000 simulated entities sets and summarized the results in terms of the Positive Predictive Value (PPV) of the applied GSEA-LFC estimator under various levels of error *ϵ*^⊥^. By design, the PPV equals 100% when the assumed value 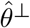 is equal to the true value 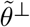 (when *ϵ*^⊥^ = 0) but may decrease when |*ϵ*^⊥^| *>* 0. Figure 1c summarizes those results and shows that the PPV can decrease to below 50% with errors on the order of *±*0.5. In words, when the average difference in scales between the two conditions is 1.65 times larger (*e*^0.5^) or 0.6 times lower (*e*^−0.5^) than assumed, more than half of the entity sets identified as significantly enriched or depleted are false positives.

Finally, in Supplementary Section 3, we expand upon these simulation studies and show how the PPV of a GSEA-LFC applied estimator varies with different LFC distributions, entity set sizes, and total number of entities. Of these factors, asymmetric LFC distributions (having more entities increase than decrease or vice versa) led to the most striking drops in PPV (to just 0.2%) with only slight error in the assumed scale (*ϵ*^⊥^ = *±*0.6). This result reinforces recent work showing the dramatic impact of compositional asymmetry on the fidelity of differential analysis [23].

### 2.4 LFC Sensitivity Analysis of Real Data

Here we expand on the results of the prior section and show that, in the context of real data, many entity sets are scale reliant. In addition, this section specifically studies those rare entity sets for which GSEA is scale invariant and develops a hypothesis test for DSA that is designed to exploit those sets.

We analyzed two previously published studies that used GSEA-LFC applied estimators to perform DSA. The first compared gene pathways expression in healthy versus normal-adjacent-totumor thyroid tissue [4]. The second compared the abundance of different microbe sets in the oral microbiota of smokers versus non-smokers [24]. We leave discussion of the microbiome study to Supplementary Section 4, as it resembles the thyroid tissue analysis and demonstrates that our conclusions hold beyond gene expression analysis. For both analyses, prior literature was used to determine upper and lower bounds for biologically plausible levels of error (*ϵ*^⊥^) in scale assumptions (see Methods).

The sensitivity of GSEA-LFC to errors in scale assumptions varied substantially over the 50 hallmark gene sets analyzed in Aran et al. [4] (Fig. 2). Some gene sets, such as the Myogenesis and KRAS Signalling Down pathways, were largely insensitive to errors in scale assumptions (they were significant over all *ϵ*^⊥^ tested). Other sets such as the MYC Targets V2 gene set were highly sensitive (significant only over a narrow range of *ϵ*^⊥^). With an error as small as *ϵ*^⊥^ = −0.1, multiple gene sets identified as enriched would no longer be significant (e.g., the columns Inflammatory Response to DNA Repair).

**Figure 2:**
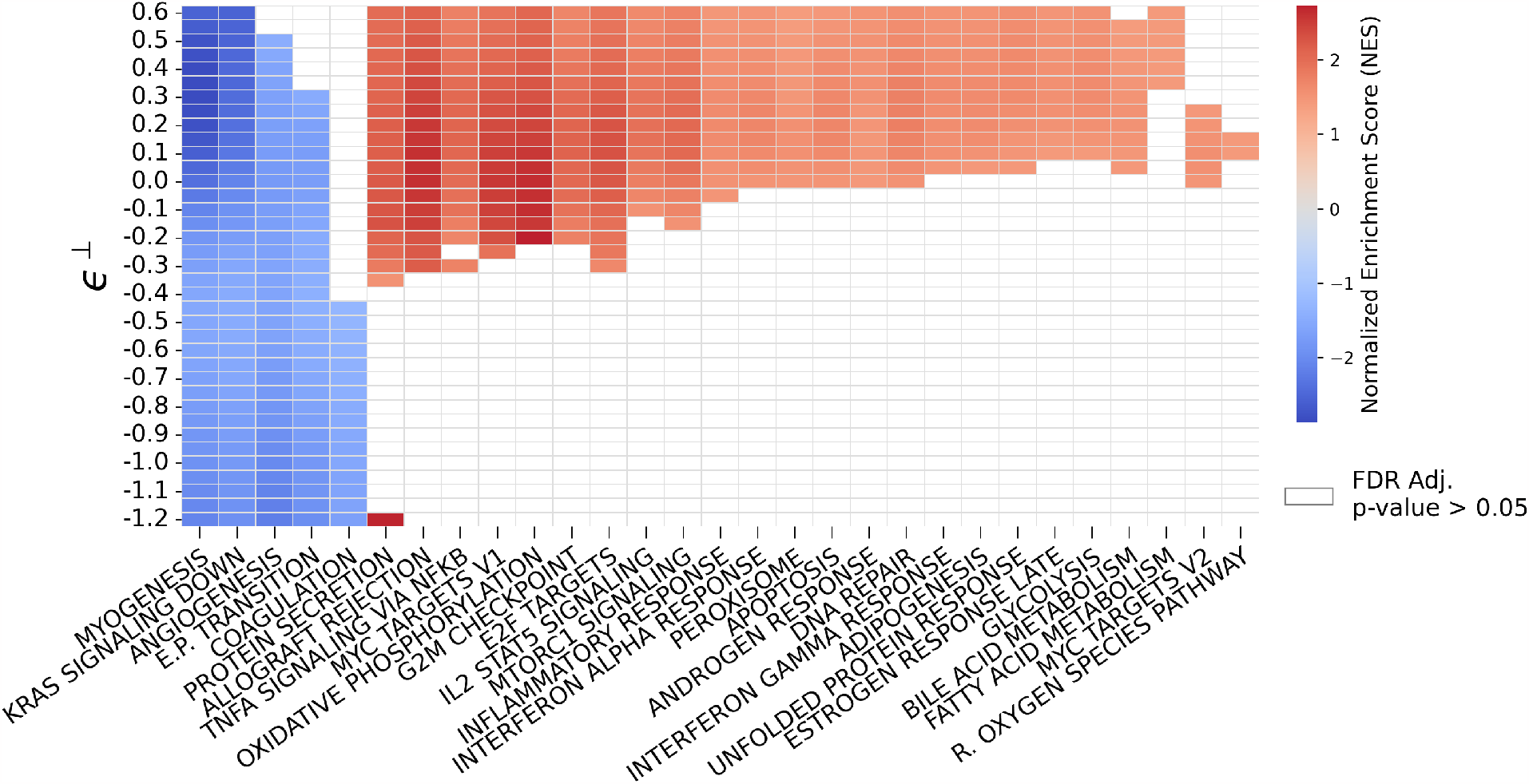
In the context of real data, GSEA-LFC applied estimators can demonstrate substantial sensitivity to errors in scale assumptions. LFC Sensitivity Analysis was used to replicate the results of Aran et al. [4], which used GSEA-LFC to compare differential pathway enrichment between normal-adjacent-to-tumor and healthy thyroid tissue. We explored sensitivity to errors in scale assumptions with the CLR assumption which, for this study, equates to 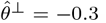. For an error *ϵ*^⊥^ (Y-axis), the implied true log-fold-change of scales is given by 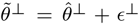. Higher (or lower) values of *ϵ*^⊥^ correspond to a higher (or lower) scale in normal-adjacent-to-tumor compared to healthy tissue than assumed. The range of *ϵ*^⊥^ ∈ [−1.2, 0.6] was informed by prior research on how much scales can vary between conditions in similar experiments; it is asymmetric to account for the CLR assumption 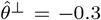(see Methods for full details). Higher values (red) of the NES correspond to more enrichment in normal-adjacent-to-tumor tissue, and lower values (blue) more enrichment in healthy tissue.

Pathways such as the Myogenesis and KRAS Signaling Down pathways, which were largely insensitive to scale errors, led us to hypothesize that there could be gene sets for which GSEA was scale invariant: where the value of 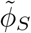 was invariant to any error *ϵ*^⊥^ ∈ (−∞, ∞). That such gene sets could exist is a remarkable feature of DSA; for DA (e.g., equation (2)), the target estimand is scale reliant regardless of the entity in question. In fact, we found such entity sets exist in both our real and simulated analyses (see Supplementary Section 5). While these entity sets are not common, their existence allowed us to develop a novel hypothesis test for DSA that provably controls Type-I error even in the presence of erroneous scale assumptions. We call this the LFC Sensitivity Analysis Test. In Supplementary Section 5 we detail this test and characterize its statistical power.

### 2.5 GSEA with Compositional Weighting

Under the SRI framework, the distinction between whether a problem is scale reliant or scale invariant depends on the scientific goal (the target estimand). Until now, we have only considered target estimators and estimands defined by replacing estimates in an applied estimator with their corresponding true values in a target estimator. For example, we defined the GSEA-LFC target estimand 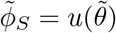 by replacing the estimates 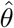 with their true values 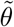 in the GSEA-LFC applied estimator 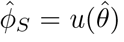. This approach allowed us to study how errors in scale assumptions, used in the estimation of *θ*, propagate into the estimation *ϕ*_*S*_. In this section, we take a different approach and assume that the applied estimator a researcher uses is tautologically consistent with their research goals. For a researcher using the GSEA-LFC applied estimator, we assume that the target estimand is 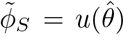. In this case, there are no scale assumptions, as there is no discrepancy between the methods applied and the goals of an analysis. Moreover, without scale assumptions, there is no need for sensitivity analysis. Instead, the challenge in this section is to interpret the research goals implied by this target estimand. This leads us to a deeper understanding of when researchers should be concerned about potential errors in scale assumptions.

To better express our meaning under this new approach, we modify the notation used in the prior sections. Rather than using 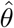 to denote estimated LFCs, where the superscript 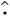 emphasized that this was an estimate, we now use the notation 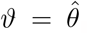. The lack of a 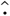 emphasizes that this is not simply an estimate or approximation but a quantity of direct scientific interest that is tautologically free of potential error. For the same reason, we replace 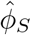 with *φ*_*S*_. For a researcher using an applied estimator of the form 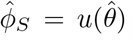, we now have a target estimand of the form 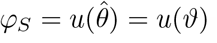. This notation also emphasizes that this target estimator is not the same as the GSEA-LFC target estimator in equation (2). For concreteness, we again focus on the common use of the CLR assumption in LFC estimation: let *ϑ* denote the log-fold change of CLR transformed abundances (equation (4)). We call *φ*_*S*_ the GSEA-CLR target estimand. How do we interpret the scientific goal represented by this estimand?

In Supplementary Section 6, we demonstrate that the LFC of CLR transformed abundances (*ϑ*_*d*_) is related to the LFC of the *d*-th entity (*θ*_*d*_) by the equation:

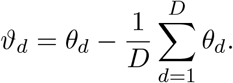

In words, researchers purposefully choosing to analyze LFCs of CLR transformed abundances (*ϑ*_*d*_) (as opposed to LFCs of actual abundances, *θ*_*d*_) do not care if an entity is truly increased or decreased between conditions, only that the entity is increased or decreased relative to the average change of all of the entities. This distinction is shown visually in Figure 3. Extending this intuition to applying GSEA to *ϑ*, researchers purposefully taking this analysis approach are only concerned about whether the entities of a set *S* are non-randomly increased or decreased between conditions *relative to the average change of all the entities*. For example, such researchers could still be interested in an entity set that is not *actually* enriched (or depleted) between conditions, but is only relatively enriched (or relatively depleted) when *compared* to the entities not in the set. Two examples will solidify this intuition and illustrate the practical implications of understanding this distinction.

**Figure 3:**
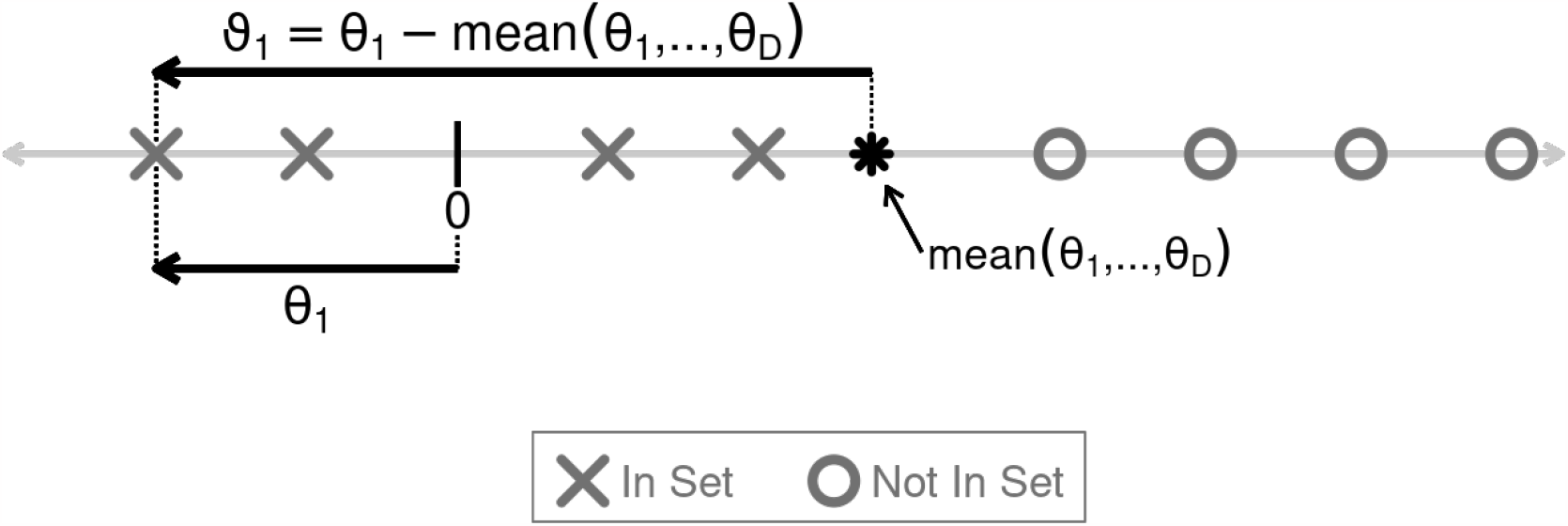
A visual depiction of the difference between Log Fold Changes (LFCs), *θ* defined by equation (2), and LFCs of CLR transformed amounts, *ϑ* defined by equation (4). The X’s and O’s are plotted on a number line and represent the LFCs of eight entities in and not in some set, *S*, of interest. In this illustration, none of the entities in the set (the X’s) are strongly increased or decreased in amount between conditions: for *d* ∈ *S, θ*_*d*_ ≈ 0. Still, each of these entities has a negative *ϑ*_*d*_, as their LFCs are each less than the mean LFC of all the entities. According to the GSEA-CLR target estimand, this set would therefore be depleted whereas it is neither depleted nor enriched according to the GSEA-LFC target estimand.

Consider two researchers, Researcher A and Researcher B. Researcher A wants to identify if a particular genomic trait confers a selective advantage in a mixed microbial community exposed to an antibiotic. Researcher A does not care if the trait actually stimulates growth in the presence of the antibiotic, only that microbes with that trait increase in abundance *relative to microbes that do not have that trait*. This goal is more consistent with the GSEA-CLR target estimand than the GSEA-LFC target estimand. As a result, Researcher A can apply GSEA to LFCs estimated using the CLR assumption without needing to consider potential error in scale assumptions. In contrast to Researcher A, Researcher B wants to identify gene pathways that are differentially activated in diseased compared to healthy tissue. Researcher B wants to understand which pathways play a causal role in the disease. Researcher B is not interested in a pathway that is unrelated to disease and is simply enriched relative to some other pathway that is repressed in disease. Researcher B’s scientific goal is more consistent with the GSEA-LFC target estimand than the GSEA-CLR target estimand. As this is a scale reliant target estimand, we would recommend that Researcher B considers potential error in scale assumptions regardless of the applied estimator chosen.

Extending beyond the GSEA-CLR target estimand, Supplementary Section 6 shows that the GSEA-CLR target estimand is actually just one of an infinite number of target estimands for DSA that are built around GSEA yet are scale invariant. We call this broader class *GSEA with Compositional Weighting* (GSEA-CW). Appealingly, each target estimand in this class can be naturally paired with a different applied estimator, which can in turn be computed from observed data. In Supplementary Section 6 we provide further characterization of this class to help researchers understand both the types of scientific goals that are invariant to erroneous scale assumptions and how to identify appropriate analytical tools given such goals. In Supplementary Section 7, we illustrate some of the advantages of this class over an alternative scale invariant approach to DSA recently proposed by Nguyen et al. [25].

## 3 Discussion

Differential Set Analysis (DSA) is a core analysis performed in modern biomedical research [26]; it is used to identify sets of entities that are differentially enriched or depleted between two experimental conditions. Although the majority of our presentation focused on the analysis of gene expression data using GSEA applied to estimated LFCs, we also showed that these same problems can be found when studying 16S rRNA microbiome data or when investigating other popular methods such as CAMERA (Supplementary Section 1). In all these cases, we arrived at the same two conclusions: 1. for common scientific goals, even slight errors in scale assumptions can lead to false positives in DSA; 2. sensitivity analyses provide a powerful tool for mitigating these problems and identifying potential false positives. Overall this work provides tools to mitigate the impact of erroneous scale assumptions, along with detailed discussions and results to illustrate the types of scientific goals that are invariant to such errors.

It is our hope that this work catalyzes future discussions and research into the role of scale limitations in DSA. We describe two areas of promising future research and refinement.

The concept of target estimands highlights how the scientific goals of a study dictate the impact of erroneous scale assumptions. In this article we have discussed a number of target estimands such as GSEA-LFC and GSEA-CW. We also discussed a CAMERA based estimand in Supplementary Section 1. Still, we expect that some researchers will feel that they perform DSA for reasons not represented in the estimands we have studied. For example, researchers may perform GSEA on LFCs but be interested in population level LFC estimates, where instead of a mean in equation (2), there is an expectation with respect to some population-level model. We believe the study of DSA from the perspective of different target estimands represents a prime area for future research.

This article has focused on DSA performed using sequence count data. Yet there has been an increasing interest in combatting scale limitations by combining sequence count data with secondary measurements designed to directly measure the system scale. For example, in the study of human microbiota, some researchers use qPCR or flow cytometry to measure the total 16S rRNA concentration or the total cellular concentration in a sample [13]. While we believe that the field’s increasing interest in these types of measurements necessitates more careful study, we suspect these measurements may be more limited than often discussed. Putting aside issues of the accuracy and precision of such measurements [11], we expect that the scale measured by these technologies may not accord with a researcher’s desired notion of amount. Many researchers use qPCR to measure the total concentration of 16S rRNA in fecal material. In the context of DSA, how often are researchers interested in identifying if the concentration of 16S rRNA from microbes in fecal material is enriched or depleted? Even if this was a concentration measurement from the colon (rather than stool), we expect there are at least some researchers interested in defining enriched or depleted sets with regards to alternative notions of amount (different definitions of scale). As a result, we expect that these technologies will not solve the problem of scale limitations for all researchers. Neither do we claim that our methods are so general as to solve the problem of scale limitations. Still, we note that our methods are not tied to a single notion of scale and may therefore have some advantages over these external measurement-based approaches.

## 4 Methods

### 4.1 Preprocessing and Differential Analysis of Thyroid Tissue RNA-seq Data

Following Aran et al. [4], we downloaded pre-processed read count data for healthy and normaladjacent-to-tumor tissue from the Genotype-Tissue Expression project (GTEx) and The Cancer Genome Atlas (TCGA) via NCBI’s Gene Expression Omnibus (GEO) under accession numbers GSE86354 and GSE62944, respectively [27, 28]. The downloaded read count data were further processed to use the same 16038 genes, 361 healthy thyroid samples, and 59 normal-adjacent-to-tumor thyroid samples used by Aran et al [4]. LFCs were estimated using the Songbird multinomial logistic-normal regression model (Version 1.0.3) [17] including an intercept term and a binary condition indicator (healthy versus normal-adjacent-to-tumor). The model was trained with default parameters and validated by ensuring cross validation error plateaued over training epochs.

### 4.2 LFC Sensitivity Analysis of Thyroid Data

Following Aran et al. [4], GSEA p-values were calculated using a list of 50 hallmark gene sets from the Molecular Signature Database (MSigDB, Version 7.4.0) [29] and FDR corrected as described in Subramanian et al. [1]. P-values were calculated using 5000 entity set label permutations.

LFC Sensitivity Analysis was performed over the range *ϵ*^⊥^ ∈ [−1.2, 0.6] in increments of 0.05. This range of *ϵ*^⊥^ was chosen based on prior literature studying tumor versus normal tissue which suggested that total RNA abundance between these conditions could vary by as much as 2.5 fold [30]. Combining this range (*±*2.5 fold) with this dataset’s CLR estimate of 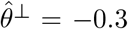 implied a range of *ϵ*^⊥^ of 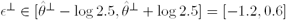.

### 4.3 Simulated LFC Sensitivity Analysis using the Positive Predictive Value

The simulated LFC Sensitivity Analysis presented in this work was summarized using Positive Predictive Value (PPV). For each value of *ϵ*^⊥^ ∈ [−2, 2] considered, PPV was calculated as the percentage of entity sets for which 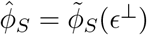 when 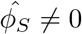 where 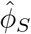 is the DSA estimate under the CLR assumption (when *ϵ*^⊥^ = 0) and 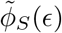 denotes the true value of *ϕ*_*S*_ when the error in the LFC estimates is equal to *ϵ*^⊥^. It follows that, by definition, the PPV is equal to 100% when *ϵ*^⊥^ = 0.

## 5 Code Availability

All code needed to reproduce the analyses in this article can be found at

https://github.com/kyle-mcgovern/DSAScaleError.

## 6 Acknowledgements

We thank Rachel Silverman and Yen Duong for their manuscript comments. J.D.S and M.P.N were supported in part by NIH 1R01GM148972-01. K.C.M was supported in part by the Computation, Bioinformatics, and Statistics (CBIOS) Training Program, NIH 5T32GM102057-10.

## 7 Contributions

K.C.M. conducted all data analyses and wrote all code. K.C.M. and J.D.S. developed the concepts of LFC Sensitivity Analysis, the LFC Sensitivity Analysis Test, and GSEA with Compositional Weighting. M.P.N. provided background knowledge needed to use Scale Reliant Interference in this work. All authors wrote and proofread the manuscript.

## Footnotes

1 This is a ratio of geometric, not arithmetic, means as this statement is made on the linear, not logarithmic, scale.

## References

1. Subramanian, A. et al. Gene set enrichment analysis: a knowledge-based approach for interpreting genome-wide expression profiles. Proc. Natl. Acad. Sci. U.S.A. 102, 15545–15550 (2005).

2. Kou, Y., Xu, X., Zhu, Z., Dai, L. & Tan, Y. Microbe-set enrichment analysis facilitates functional interpretation of microbiome profiling data. Sci. Rep. 10, 1–12 (2020).

3. Verfaillie, A. et al. Decoding the regulatory landscape of melanoma reveals TEADS as regulators of the invasive cell state. Nat. Commun. 6, 6683 (2015).

4. Aran, D. et al. Comprehensive analysis of normal adjacent to tumor transcriptomes. Nat. Commun. 8, 1–13 (2017).

5. Murohashi, M. et al. Gene set enrichment analysis provides insight into novel signalling pathways in breast cancer stem cells. Br. J. Cancer 102, 206–212 (2010).

6. Props, R. et al. Absolute quantification of microbial taxon abundances. ISME J. 11, 584–587 (2017).

7. Gloor, G. B., Macklaim, J. M., Pawlowsky-Glahn, V. & Egozcue, J. J. Microbiome datasets are compositional: And this is not optional. Front. Microbiol. 8, 1–6 (2017).

8. Robinson, M. D. & Oshlack, A. A scaling normalization method for differential expression analysis of RNA-seq data. Genome Biol. 11, R25 (2010).

9. Anders, S. & Huber, W. Differential expression analysis for sequence count data. Genome Biol. 11 (2010).

10. Quinn, T. P., Erb, I., Richardson, M. F. & Crowley, T. M. Understanding sequencing data as compositions: An outlook and review. Bioinformatics 34, 2870–2878 (2018).

11. Nixon, M. P., Letourneau, J., David, L., Mukherjee, S. & Silverman, J. D. Scale Reliant Inference. Preprint at http://arxiv.org/abs/2201.03616 (2022).

12. Aitchison, J. Principles of Compositional Data Analysis. Lect. Notes Monogr. Ser. 24, 73–81 (1994).

13. Jian, C., Luukkonen, P., Yki-Järvinen, H., Salonen, A. & Korpela, K. Quantitative PCR provides a simple and accessible method for quantitative microbiota profiling. PLoS ONE 15, 1–10 (2020).

14. Vandeputte, D. et al. Quantitative microbiome profiling links gut community variation to microbial load. Nature 551, 507–511 (2017).

15. Silverman, J. D. et al. Measuring and mitigating PCR bias in microbiota datasets. PLoS Comput. Biol. 17, 1–17 (2021).

16. Fernandes, A. D. et al. Unifying the analysis of high-throughput sequencing datasets: Characterizing RNA-seq, 16S rRNA gene sequencing and selective growth experiments by compositional data analysis. Microbiome 2, 1–13 (2014).

17. Morton, J. T. et al. Establishing microbial composition measurement standards with reference frames. Nat. Commun. 10 (2019).

18. Love, M. I., Huber, W. & Anders, S. Moderated estimation of fold change and dispersion for RNA-seq data with DESeq2. Genome Biol. 15, 1–21 (2014).

19. Wiebe, D. S. et al. Fold-Change-Specific Enrichment Analysis (FSEA): Quantification of Transcriptional Response Magnitude for Functional Gene Groups. Genes 11, 434 (Apr. 2020).

20. Wu, D. & Smyth, G. K. Camera: A competitive gene set test accounting for inter-gene correlation. Nucleic Acids Res. 40, 1–12 (2012).

21. Gatti, D. M., Barry, W. T., Nobel, A. B., Rusyn, I. & Wright, F. A. Heading down the wrong pathway: on the influence of correlation within gene sets. BMC Genomics 11, 574–574 (2010).

22. Tamayo, P., Steinhardt, G., Liberzon, A. & Mesirov, J. P. The limitations of simple gene set enrichment analysis assuming gene independence. Stat. Methods Med. Res. 25, 472–487 (2016).

23. Wu, J. R., Macklaim, J. M., Genge, B. L. & Gloor, G. B. in Advances in Compositional Data Analysis: Festschrift in Honour of Vera Pawlowsky-Glahn 329–346 (Springer International Publishing, Cham, 2021).

24. Beghini, F. et al. Tobacco exposure associated with oral microbiota oxygen utilization in the New York City Health and Nutrition Examination Study. Ann. Epidemiol. 34, 18–25.e3 (2019).

25. Nguyen, Q. P., Hoen, A. G. & Frost, H. R. CBEA: Competitive balances for taxonomic enrichment analysis. PLoS Comput. Biol. 18, 1–24 (2022).

26. Maleki, F., Ovens, K., Hogan, D. J. & Kusalik, A. J. Gene Set Analysis: Challenges, Opportunities, and Future Research. Front. Genet. 11, 1–16 (2020).

27. Lonsdale, J. et al. The Genotype-Tissue Expression (GTEx) project. Nat. Genet. 45, 580–585 (2013).

28. Rahman, M. et al. Alternative preprocessing of RNA-Sequencing data in the Cancer Genome Atlas leads to improved analysis results. Bioinformatics 31, 3666–3672 (2015).

29. Liberzon, A. et al. The Molecular Signatures Database Hallmark Gene Set Collection. Cell Syst. 1, 417–425 (2015).

30. Lin, C. Y. et al. Transcriptional amplification in tumor cells with elevated c-Myc. Cell 151, 56–67 (2012).

