## Supplemental Text and Figures for "Addressing Erroneous Scale Assumptions in Microbe and Gene Set Enrichment Analysis"

#### 1 Inter-Entity Correlations

In the main text we focus on GSEA-LFC with respect to entity-label permutations which implicitly assumes that the entities are statistically independent: an assumption that can be violated in practice leading to false positives [1–3]. Here we extend the results of the main text to three DSA methods that account for inter-entity correlations. The first two methods are GSEA-LFC-S and GSEA-CLR-S. These methods are GSEA-LFC and GSEA-CLR respectively but with the suffix *-S* indicating the use of sample-label (rather than entity-label) permutations which maintain the correlation structure of the data [4, 5]. We also consider the CAMERA [2] method which looks for a difference in the mean LFC between entities in and out of the set using a t-test while including an inter-entity correlation correction factor.

We initially explored the sensitivity of GSEA-LFC-S to errors in scale assumptions using the Aran et al. thyroid tissue dataset [6] that we reanalyzed in Section 2.4 of the main text. Remarkably, in changing from using entity-label permutations (as used by Aran et al. [6]), to sample-label permutations, GSEA-LFC-S returned no significant hits. This suggests that the results reported in Aran et al. could be due entirely to false positives secondary to an erroneous assumption that the entities were independent (i.e., the gene expression was not correlated). While these results highlight the importance of considering inter-entity correlations, it also implied that this dataset was likely a poor test-case for studying the role of scale-limitations when accounting for inter-entity correlations. As a result, here we focus on a different dataset consisting of 184 samples of either healthy or cancerous breast tissue and a much larger list of 4622 genes sets (see Supplementary Section 1.2).

Performing a sensitivity analysis of this breast cancer dataset also required abandoning LFC Sensitivity Analysis in favor of Scale Sensitivity Analysis [7]. For LFC Sensitivity Analysis we performed a sensitivity analysis of the target estimand ($\tilde{\phi}$) as a function of error $\epsilon^{\perp}$ in the estimated LFCs ($\hat{\theta}$): $\tilde{\phi}_S = u(\hat{\theta} + \epsilon^{\perp})$. The simplicity of this approach relied in part on the fact that the estimate of the log-fold-change in scales between conditions (i.e., $\hat{\theta}^{\perp}$) did not change under entity-label permutations. However, for GSEA-LFC-S the estimate $\hat{\theta}^{\perp} = \text{mean}_{n:x_n=1}(\log W_n^{\perp}) - \text{mean}_{n:x_n=0}(\log W_n^{\perp})$, will change depending on which samples are assigned to the healthy and diseased groups after sample-label permutation (i.e., whether $x_n = 1$ or $x_n = 0$ for each sample $n$ after shuffling the binary sample labels). Instead, we perform Scale Sensitivity Analysis by evaluating the error in the applied estimand after adding sample-specific scale error $\alpha_n^{\perp}$ to each sample on the log scale. As in LFC Sensitivity Analysis, we assume that there is limited compositional error such that $\tilde{W}^{\parallel} \approx \hat{W}^{\parallel}$. Similar to LFC Sensitivity Analysis we may then express the estimated scaled system on the log scale for taxa $d$ and sample $n$ as

$$\begin{aligned}
\log \tilde{W}_{dn} &= \log \tilde{W}_{dn}^{\parallel} + \log \tilde{W}_n^{\perp} && (1) \\
&\approx \log \hat{W}_{dn}^{\parallel} + \left(\log \hat{W}_n^{\perp} + \alpha_n^{\perp}\right) && (2) \\
&\approx \log \hat{W}_{dn} + \alpha_n^{\perp}. && (3)
\end{aligned}$$

Scale Sensitivity Analysis is a sensitivity analysis comparing the applied estimand $\hat{\phi}$ to the target estimand $\tilde{\phi}$ when the applied estimand has errors $\alpha_1^{\perp}, \ldots, \alpha_n^{\perp}$: $\tilde{\phi}_S = u\left(f\left(\log \hat{W}_{\cdot,1} + \alpha_1, \ldots, \log \hat{W}_{\cdot,n} + \alpha_n^{\perp}\right)\right)$. The function $f$ returns LFCs $\hat{\theta}$ for GSEA-LFC-S and GSEA-CLR-S. For CAMERA the matrix $\hat{W}$ is used directly, without first summarising the data as LFCs, so $f$ can be thought of as the identity function.

We conducted two Scale Sensitivity Analyses whose results are visualized in Figure S1. In the first sensitivity analysis shown in Figure S1a we simulate error $\alpha_n^{\perp}$ as some constant $\delta^{\perp}$:

$$\alpha_n^{\perp} = \begin{cases} \delta^{\perp}, & \text{if } x_n = 1 \\ 1, & \text{if } x_n = 0. \end{cases}$$

In this particular sensitivity analysis, error ($\delta^{\perp}$) is constant across samples in the diseased condition and 0 for samples in the healthy experimental condition. As a result this analysis has a similar interpretation to error in $\hat{\theta}^{\perp}$ (i.e., $\tilde{\theta}^{\perp} = \hat{\theta}^{\perp} + \epsilon^{\perp}$) because

$$\begin{aligned}
\tilde{\theta}^{\perp} &= \underset{n:x_n=1}{\text{mean}}(\log \hat{W}_n^{\perp} + \delta^{\perp}) - \underset{n:x_n=0}{\text{mean}}(\log \hat{W}_n^{\perp} + 0) \\
&= \underset{n:x_n=1}{\text{mean}}(\log \hat{W}_n^{\perp}) - \underset{n:x_n=0}{\text{mean}}(\log \hat{W}_n^{\perp}) + \delta^{\perp} \\
&= \hat{\theta}^{\perp} + \delta^{\perp}.
\end{aligned}$$

It should be noted however that Scale Sensitivity Analysis is still required for this analysis as the diseased and healthy condition labels ($x_n$) change with sample-label shuffling, and thus the estimated means change as well.

In the second Scale Sensitivity Analysis show in in Figure S1b we simulated distinct, independent error in each $\log \hat{W}_n^{\perp}$ for some constant $\gamma^{\perp}$ from the uniform distribution (denoted as $U$):

$$\alpha_n^{\perp} \sim U[-\gamma^{\perp}, \gamma^{\perp}].$$

Unlike the first Scale Sensitivity Analysis, here an element of randomness has been introduced in sampling from the uniform distribution and thus we performed this Scale Sensitivity Analysis for 50 independent samples of $\alpha_1^{\perp}, ..., \alpha_N^{\perp}$. We discuss the results of these two Scale Sensitivity Analyses in the remainder of this section.

Like the LFC Sensitivity Analyses in the main text, here we use the Positive Predictive Value (PPV) as a measure of sensitivity for different values of $\alpha_1^{\perp}, \ldots, \alpha_N^{\perp}$ tested. When there is no sensitivity to scale errors we expect the PPV to be 100% across all scale assumption errors tested. Our results show that GSEA-CLR-S is completely insensitive (with a PPV of 100%) to all errors in scale assumptions (Fig. S1, blue diamonds and blue box plots). GSEA-CLR-S was still able to identify 595 putative hits despite this scale insensitivity. We found that GSEA-LFC-S is sensitive to errors $\delta^{\perp}$ in $\theta^{\perp}$ (Fig. S1a, orange X symbols) with the PPV reaching as low as 54%. GSEA-LFC-S is however not nearly as sensitive to random errors in $\hat{W}^{\perp}$ (Fig. S1b, orange boxplots) with the PPV only dropping to 96%. We found that CAMERA is completely insensitive to errors in $\hat{\theta}^{\perp}$ (Fig. S1a, blue diamonds), but shows a high degree of sensitivity to errors in $\hat{W}^{\perp}$. For errors in $\hat{W}^{\perp}$ sampled uniformly when $\gamma^{\perp} = 0.05$, CAMERA's PPV dropped to 94% and to as low as 53% when $\gamma^{\perp} = 0.25$.

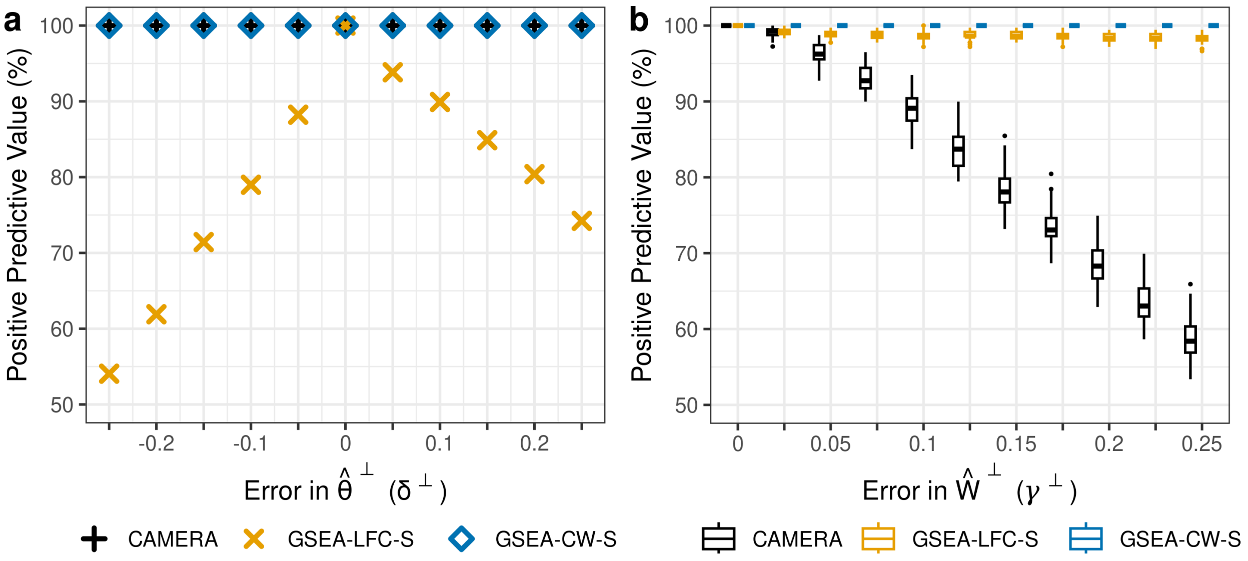


**Figure S1: Common methods that account for inter-entity correlations such as CAMERA and GSEA-LFC-S are scale reliant while GSEA-CW-S is scale invariant.** The figure represents the output of two Scale Sensitivity Analyses. Two types of errors in scale assumptions were considered: errors $\delta^{\perp}$ in the assumed expected log-fold-change of scales ($\tilde{\theta}^{\perp} = \hat{\theta}^{\perp} + \delta^{\perp}$; **Panel a**) and errors $\alpha_n^{\perp}$ in the assumed scale of individual participants ($\log \tilde{W}_n^{\perp} = \log \hat{W}_n^{\perp} + \alpha_n^{\perp}$; **Panel b**). As the latter errors are multi-dimensional they were randomly simulated as $\alpha_n^{\perp} \sim U(-\gamma^{\perp}, \gamma^{\perp})$ where $U$ is the uniform distribution. The boxplots represent the positive predictive values for 50 independent simulations at each value of $\gamma^{\perp}$. For both types of error, sensitivity was summarized for a study comparing gene expression in healthy (n=92) versus tumor (n=92) breast tissue using a database of 4622 gene sets (see Supplementary Sections 1.1 and 1.2 for further details).

We also found that the severity of the scale sensitivity for GSEA-LFC-S is gene set dependent and that this may have implications for actual research. The Cell Cycle and Mismatch Repair pathways have both been identified in the literature as enriched in breast cancer tissue [8], yet here we find that only the former pathway is enriched in breast cancer tissue across a wide range of tested values of $\delta^{\perp}$ while the latter is enriched only over a very narrow range of values of $\delta^{\perp}$ (Fig. S2).

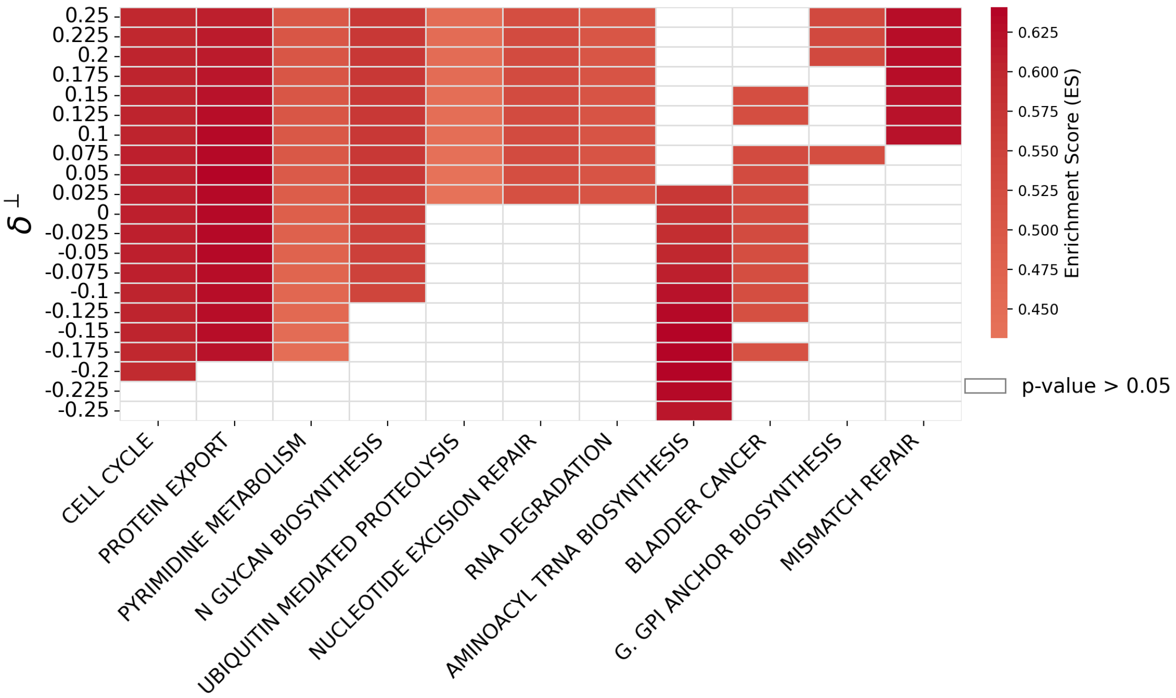


**Figure S2: Gene sets identified by GSEA-LFC-S have different degrees of sensitivity to errors in scale assumptions.** Gene set specific results are presented for the Scale Sensitivity Analysis presented in Figure S1a. $\delta^{\perp}$ may be interpreted as error in the assumed LFC in the assumed log-fold-change in scales $\hat{\theta}^{\perp}$: $\tilde{\theta}^{\perp} = \hat{\theta}^{\perp} + \delta^{\perp}$.

Overall, our results are in line with the central findings of the main text: both GSEA-LFC-S and CAMERA are sensitive to errors in scale assumptions, albeit with the added complexity that the sensitivity depends on whether there is random scale error for each independent sample (e.g., $\alpha_n^{\perp} \sim U[-\gamma^{\perp}, \gamma^{\perp}]$) or constant scale error that differs only between experimental conditions (e.g., $\delta^{\perp}$). The observation that GSEA-CLR-S is scale invariant in both of our sensitivity studies similarly aligns with our findings presented in Supplementary Section 6 that GSEA-CW (and by extension GSEA-CLR-S) is scale invariant.

#### 1.1 Methods for Preprocessing of Breast Tissue RNA-seq Data

The pre-processed read count matrices were downloaded from NCBI's GEO under accession numbers GSE86354 (for healthy tissue) and GSE62944 (for tumor tissue) [6, 9, 10]. These datasets contained samples from a variety of tissues, therefore samples were filtered until only 1,119 breast tumor tissue samples and 92 healthy breast tissue samples remained. To prevent the Scale Sensitivity Analyses from being impacted by class imbalance, we subsampled the number of tumor samples to 92 to match the number of samples in the healthy condition then merged the read count matrices together by gene name. After sample filtering, many genes contained exclusively zero counts or only a few counts in one condition. To ensure that they had some degree of signal in both conditions we only included genes if they had non-zero counts in at least 3 replicates in both the healthy and tumor conditions, resulting in a filtered set of 22,280 genes.

##### 1.2 Methods for Scale Sensitivity Analyses

Both the Scale Sensitivity Analyses were performed using the MSigDB C2 (version 7.4.0) curated list of gene sets [4, 11]. Based on the default parameters of the GSEA software package [4], only gene sets that containing between 15 and 500 genes were retained in our analysis resulting in a set of 4622 candidate gene sets.

A pseudo-count of 0.1 was added to the read count matrix to ensure stability of the CLR transform. Following the addition of a pseudo-count, scale error $\alpha_n$ was added to the columns of $\hat{W}^{CLR}$: the matrix $\hat{W}$ calculated using the CLR assumption. In the first Scale Sensitivity Analysis presented, errors $\alpha_n^{\perp} = \delta^{\perp}$ for a constant $\delta^{\perp}$ were added only to columns representing the breast tumor samples before sample-label permutation. In the second Scale Sensitivity Analysis presented, errors $\alpha_n^{\perp}$, sampled from the uniform distribution $\alpha_n^{\perp} \sim U[-\gamma^{\perp}, \gamma^{\perp}]$ for a constant $\gamma^{\perp}$, were added to all columns before sample-label permutation.

To perform GSEA-CLR-S, the matrix $\hat{W}^{CLR}$ with error added was used (after permuting the sample-labels in the case of the null distribution) to calculate LFCs $\theta$. These $\theta$ were used to calculate the GSEA-CLR ranking statistics $\vartheta_d = \theta_d - \frac{1}{D}\sum_{i=1}^{D} \theta_i$.

For CAMERA, the matrix $\hat{W}^{CLR}$ with error added was used directly. Inter-entity correlations were estimated using the *interGeneCorrelation* function in the Limma R package version 5.50.3 [12]. The inter-entity correlations were then used as input to the *cameraPR* function which returned p-values for enrichment or depletion.

All p-values for both sensitivity analyses were calculated using a significance threshold of $\alpha < 0.05$ and 5000 sample-label permutations.

#### 2 Gene Set Enrichment Analysis

In this section we provide a thorough definition of Gene Set Enrichment Analysis (GSEA) [4]. Subramanian et al. [4] define the GSEA Enrichment Score (ES) as the supremum (i.e., maximum distance from 0) of the difference of two running sums ($P_{hit} - P_{miss}$) [4]. For an entity set $S$ of size $D_S$, for each entity $d \in [1, \ldots, D]$ in the list ranked by a ranking stat $R_d$, $P_{hit}$ and $P_{miss}$ are calculated as

$$P_{hit}(S,d) = \sum_{\substack{j \leq d \\ j \in S}} \frac{|R_j|}{N_R}, \text{ where } N_R = \sum_{j \in S} |R_j|$$

$$P_{miss}(S,d) = \sum_{\substack{j \leq d \\ j \notin S}} \frac{1}{D - D_S}.$$

Letting $R_d = \theta_d$ (where $\theta_d$ represents the LFC for entity $d$) we obtain GSEA-LFC(-S). In contrast letting $R_d = f(l_d(\theta))$, where $l_d(\theta)$ denotes a log-contrast of the vector $\theta$ and $f$ represents an arbitrary real-valued function, results in CSEA-CW(-S) (see Supplementary Section 6 for more details).

A p-value is determined for GSEA using a null distribution created by permuting the entity set labels (i.e., which values of $1, \ldots, D$ are in the set $S$). If the unpermuted ES is a positive value then the p-value is calculated as the proportion of positively-valued, permuted ES that are equal to or larger than the unpermuted ES. If the unpermuted ES is a negative value it is the proportion of negatively-valued, permuted ES values that are equal to or less than the unpermuted ES. For GSEA-LFC-S and GSEA-CW-S the sample labels are permuted before calculating the ranking statistic (i.e., the binary vector $X = x_1, \ldots, x_n$ indicating which participants $n$ belong to the healthy or diseased tissue condition is permuted before calculating $R_d$), and the entity set labels are not permuted at all. P-values for GSEA using sample-label permutations are then calculated using the same process as GSEA using entity-label permutations.

### 3 A Larger Set of LFC Sensitivity Simulations for GSEA-LFC

In Section 2.3 and Figure 1c in the main text we presented a single, small simulation experiment to demonstrate that GSEA-LFC is scale reliant and that Positive Predictive Values (PPVs) can be as low as 10%. Here we describe a larger study consisting of many simulated datasets to explore how the sensitivity of GSEA-LFC to errors in scale assumptions depends on a range of factors such as the total number of entities ($D$), the size of the entity sets ($D_S$), and the distribution of the simulated LFCs ($p$).

Based on prior work (e.g., [2]), we hypothesized that highly skewed LFC distributions would result in lower PPVs. We chose the distribution of LFCs ($p$) to be either a normal distribution, uniform distribution, or left- or right-skewed distribution (Fig. S3a). The left- and right-skewed distributions were created by randomly simulating 10% of the LFC from a uniform distribution on the interval $[-1, 0]$ and $[0, 1]$, respectively, and setting the remaining 90% of LFC to 0.01 or -0.01, respectively (Fig. S3a). We chose $D$ to be 500, 10,000, or 20,000 and $D_S$ to be 15, 50, or 150. For each possible combination of $p$, $D$, and $D_S$ we simulated 10,000 entity sets and performed our PPV based LFC Sensitivity Analysis over a range of values $\epsilon^{\perp} \in (-2, 2)$ using a significance cutoff of $\alpha < 0.05$ and 5000 entity-label permutations. As hypothesized, the asymmetric LFC distributions resulted in substantially lower PPVs than the other distributions while the impact of changing $D_S$ or $D$ was minimal comparatively (Fig. S3b). These effects can be dramatic: when $D_S$=150 with an asymmetric distribution of LFCs we observed a PPV of only 0.2% with a value of $\epsilon^{\perp}$ as small as $\pm 0.6$.

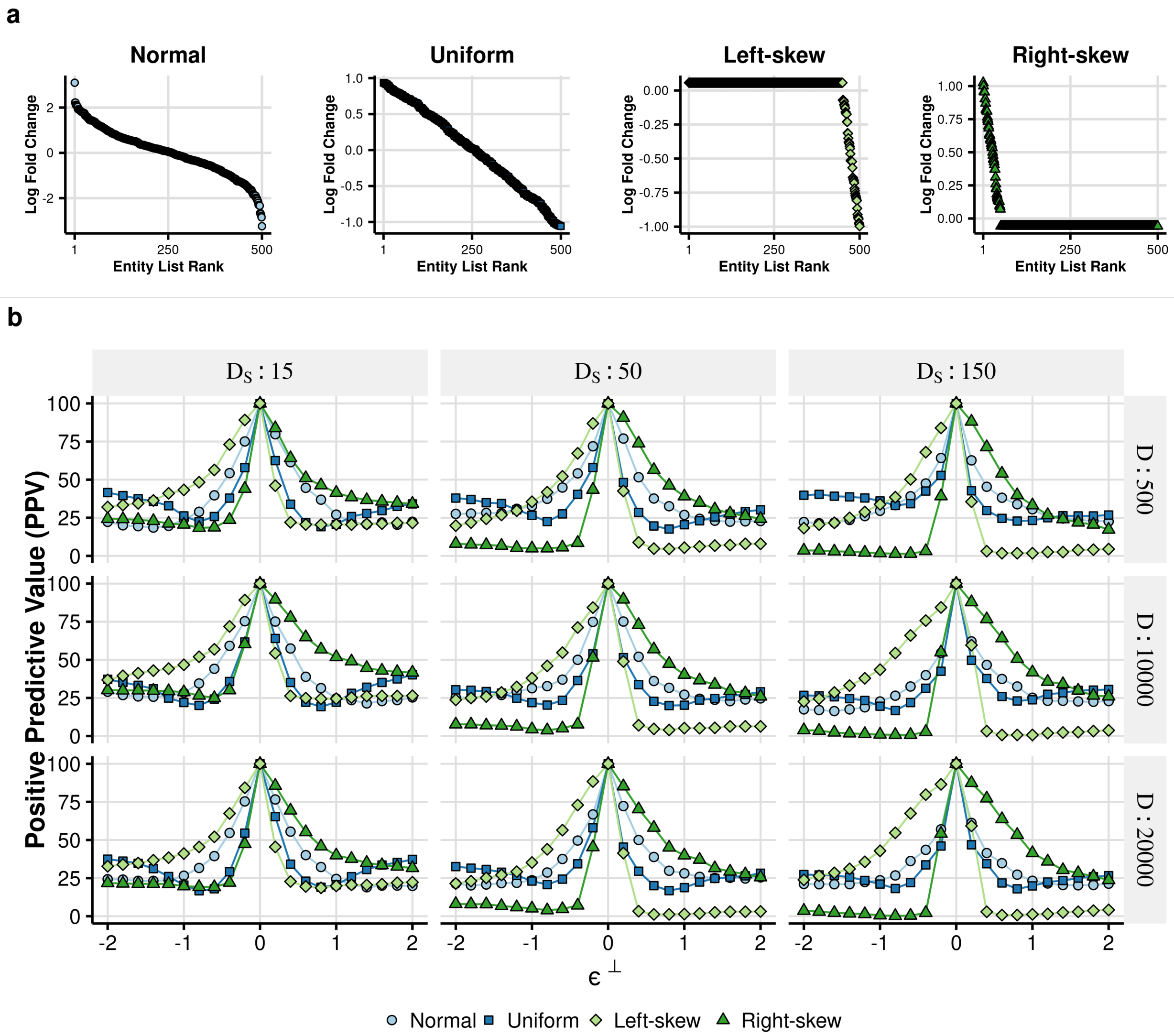


**Figure S3: A PPV based LFC Sensitivity Analysis of simulated data demonstrates that the distribution of LFC most affects the degree to which GSEA-LFC is scale reliant. (a)** Example plots of the four different distribution types that were simulated for the LFC Sensitivity Analysis. **(b)** The results of the LFC Sensitivity Analysis are plotted in a grid. The columns of the grid represent different entity set sizes ($D_S$) and the rows represent different total numbers of entities simulated ($D$). The PPV is calculated as the proportion of entity sets identified as significant ($\alpha < 0.05$) by GSEA-LFC at the assumed $\hat{\theta}^{\perp}$ (i.e., when $\epsilon^{\perp} = 0$) that were also identified when the true log-fold-change in scales is instead $\tilde{\theta}^{\perp} = \hat{\theta}^{\perp} + \epsilon^{\perp}$.

### 4 LFC and Scale Sensitivity Analyses of a Real Microbiome Dataset

In Section 2.4 and Figure 2 in the main text we presented a reanalysis of a real RNA-seq experiment using LFC Sensitivity Analysis. Here we present a reanalysis of a real 16S rRNA-seq experiment using LFC Sensitivity Analysis on real microbe sets. These results show that the findings presented in the main text for RNA-seq data may be extended to a microbe set enrichment analysis. We also present a Scale

Sensitivity Analysis to extend the results of Supplementary Section 1, which explored the sensitivity of GSEA-LFC with sample-label permutations (i.e., GSEA-LFC-S), to 16S rRNA-seq data. Specifically, we find that using GSEA-LFC(-S) does not change our conclusion that GSEA-LFC(-S) is sensitive to errors in scale assumptions, and that this sensitivity is entity-set specific.

These sensitivity analyses were performed using data from the NYC Health and Nutrition Examination Study (NYC HANES). This study compared the oral microbiomes of smokers ($n = 86$) and non-smokers ($n = 43$; see Supplementary Section 4.1). Three microbe sets were used corresponding to aerobic, anaerobic and facultative anaerobic microbes. Applying GSEA-LFC to the dataset with scale error ($\epsilon^{\perp}$) ranging from $[-1, 1]$ we find, similar to the RNA-seq study, that different microbe sets demonstrate varying degrees of scale sensitivity (Fig. 4a). The aerobic microbe set is largely insensitive to scale assumptions, as it is significantly enriched across all values of $\epsilon^{\perp}$ tested, albeit with changing enrichment scores. The facultative anaerobic (F. Anaerobic) microbe set, on the other hand, was highly sensitive as it was only enriched at a single value of $\epsilon^{\perp}$.

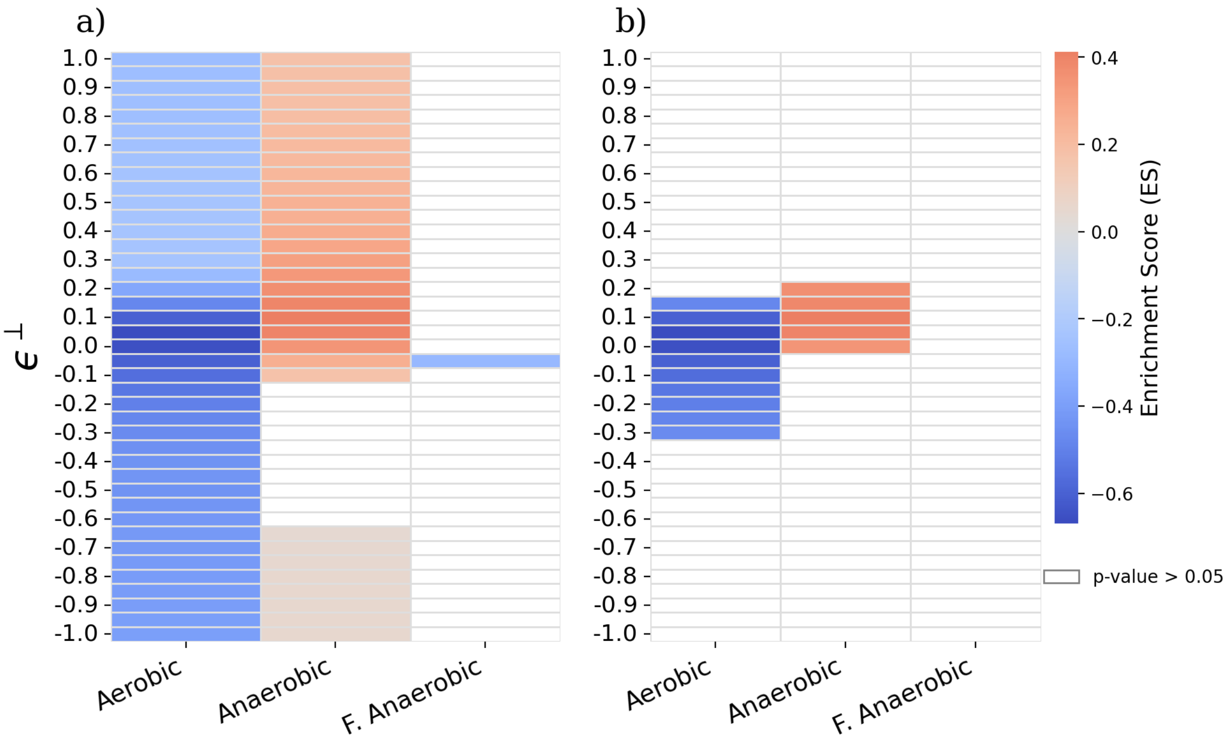


**Figure S4: Both GSEA-LFC and GSEA-LFC-S are sensitive to scale assumptions in the context of microbe set enrichment analysis.** **(a)** LFC Sensitivity Analysis using GSEA-LFC. **(b)** Scale Sensitivity Analysis using GSEA-LFC-S. Errors $\epsilon^{\perp}$ in the CLR scale assumption are shown on the Y-axis in both plots. Higher (or lower) values of $\epsilon^{\perp}$ correspond to a higher (or lower) scale in smoker compared to non-smoker samples than assumed. Higher values (red) of the Enrichment Score (ES) indicate enrichment in smokers, while lower values (blue) correspond to enrichment in non-smokers.

We see similar results for our Scale Sensitivity Analysis using GSEA-LFC-S. For this analysis $\epsilon^{\perp}$ represents constant error added to each sample derived from smokers before sample-label permutations (see Supplementary Section 4.1). Our findings are similar to the Scale Sensitivity Analysis of GSEA-LFC-S presented in Supplementary Section 1: not only was scale sensitivity observed but it was microbe-set dependent. The aerobic microbe set was less sensitive (i.e., returned more significant results over a larger range of $\epsilon^{\perp}$) than the anaerobic microbe set (Fig. 4b).

Together the results of these two sensitivity analyses reaffirm that scale sensitivity is a concern not just for gene set enrichment analysis using real RNA-seq data but for enrichment analyses of microbe sets using real 16S rRNA-seq data as well.

##### 4.1 Methods for 16S rRNA-seq LFC and Scale Sensitivity Analyses

Sensitivity analyses were performed using raw read counts from the NYC HANES 16S rRNA-seq oral microbiome dataset [13] comparing the oral microbiomes of non-smokers (n=43) to smokers (n=86) that were processed as described by Beghini et al. [14].

Three microbe sets representing aerobic, anaerobic, and facultative anaerobic microbes were used in the analysis [14]. For all analyses, LFCs were calculated after adding a pseudo-count of 0.1 to the CLR transformed read counts using equation (4) in the main text, which meant calculating $\hat{\theta}$ using the CLR assumption. The error range $\epsilon^{\perp} \in [-1, 1]$ was chosen for symmetry and to illustrate sensitivity to scale assumptions, not for addressing any specific biological questions or hypotheses.

For the Scale Sensitivity Analysis, error $\epsilon^{\perp}$ was added to counts after adding a pseudo-count and applying the CLR transform to the counts. This error $\epsilon^{\perp}$ was added only to samples derived from smokers condition, no error was added to samples derived from non-smokers. After adding the error $\epsilon^{\perp}$, p-values were calculated using GSEA [4] with 5000 sample-label permutations, which we refer to as GSEA-LFC-S.

#### 5 The LFC Sensitivity Analysis Test

Section 2.4 of the main text identified two gene sets (Myogenesis and KRAS Signalling Down) that were completely insensitive to scale errors over the range of tested error levels $\epsilon^{\perp}$. This finding led us to wonder if, for a particular entity set $S$, the GSEA-LFC target estimand could actually be scale invariant: where the value of $\tilde{\phi}_S$ was fixed at either 1 or $-1$ over the entire range $\epsilon^{\perp} \in (-\infty, \infty)$. Based on the findings of Section 2.4 in the main text, it is clear that this must not be common in practice. Yet, remarkably, we find that such gene sets can sometimes be identified in practice. This suggests that, for at least a subset of real gene sets, we can develop a rigorous hypothesis test for GSEA-LFC that provably controls Type-I error regardless of errors in scale assumptions while simultaneously having non-zero statistical power. We present such a test here.

Let $p_{\epsilon^{\perp}}$ denote the GSEA-LFC p-value at a particular value of $\epsilon^{\perp}$. We propose a new hypothesis test which we call the LFC Sensitivity Analysis Test, defined by the composite p-value $p = \max_{\epsilon^{\perp} \in (-\infty,\infty)} p_{\epsilon^{\perp}}$.

In words, this new test only rejects the null hypothesis if the GSEA-LFC p-value is less than some threshold (e.g., $p_{\epsilon^\perp} < 0.05$) over all possible errors $\epsilon^\perp$. Notably, prior work on statistical inference in the presence of nuisance parameters proves that this test rigorously controls Type-I error [15].

We explored the statistical power of the LFC Sensitivity Analysis Test in the context of real data by applying it to the same thyroid tissue dataset analyzed in Section 2.4 of the main text. Calculating statistical power (i.e., the probability of detecting a true positive) typically requires simulation studies in order to know which gene sets are truly enriched or depleted. As we lack validated approaches for simulating datasets with known set enrichment we instead used LFC Sensitivity Analysis to calculate statistical power as a function of potential error in scale assumptions using the thyroid dataset along with a set of 4441 gene sets (see methods in Supplementary Section 5.1). The results of this power analysis of the LFC Sensitivity Analysis Test are shown in Figure S5. We find that the average statistical power for the LFC Sensitivity Analysis Test is likely in the range of 6% to 13.5%. Out of the 4441 gene sets tested, the LFC Sensitivity Analysis Test identified 96 entity sets as significantly enriched or depleted. These results show that the LFC Sensitivity Analysis Test exhibits non-zero power and thus may have practical utility in research.

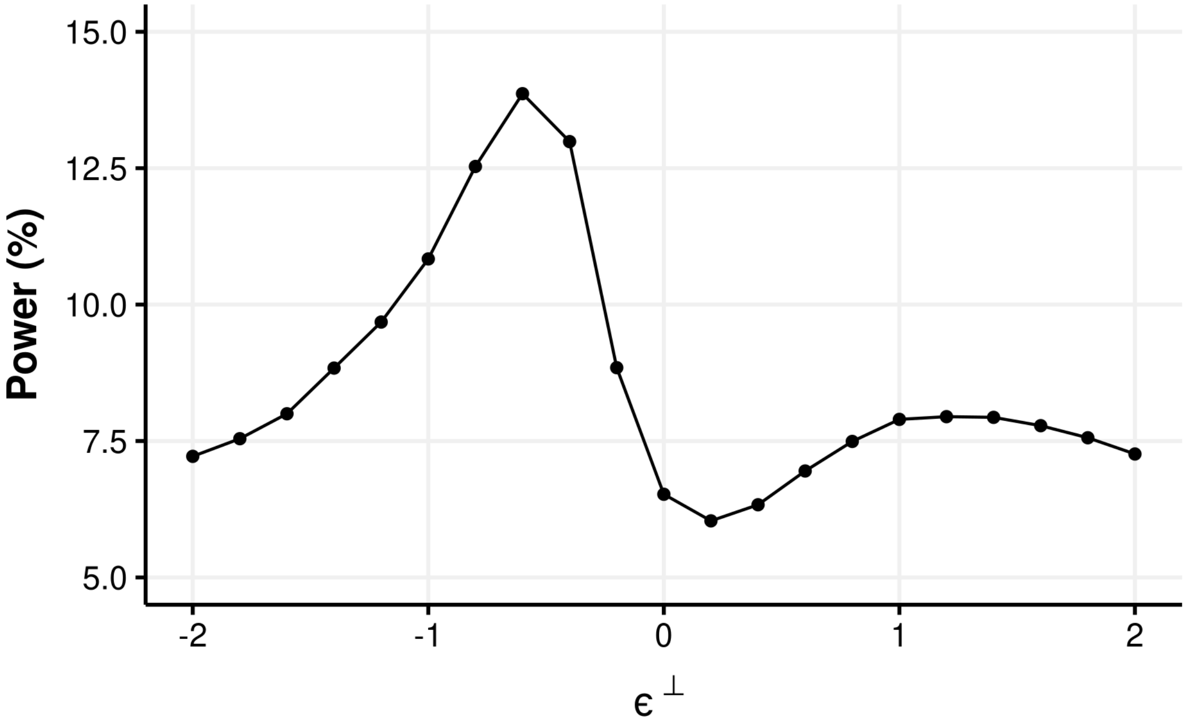


**Figure S5: The LFC Sensitivity Analysis Test has non-zero power.** The thyroid normal-adjacent-to-tumor versus healthy tissue data set described in Section 4.1 in the main text was used along with 4441 gene sets to calculate power. Power is calculated as the percentage of gene sets identified as significant with GSEA-LFC when $\tilde{\theta} = \hat{\theta} + \epsilon^{\perp}$ that were also identified as significant with the LFC Sensitivity Analysis Test using a significance of $\alpha < 0.05$.

#### 5.1 Methods

Power was estimated using LFCs estimated from the thyroid normal-adjacent-to-tumor versus healthy study as described in Section 4.1 in the main text and using the MSigDB C2 (version 7.4.0) curated list of gene sets [4, 11]. Based on the default parameters of the GSEA software package [4], only gene sets that containing between 15 and 500 genes were retained in our analysis resulting in a set of 4441 candidate gene sets. Power is calculated at any single value of $\epsilon^{\perp}$ as the percentage of gene sets that GSEA-LFC identified as significant that LFC Sensitivity Testing (using GSEA-LFC) also identified as significant. P-valus for the LFC Sensitivity Analysis Test (e.g., $p = \max_{\epsilon^{\perp}} p_{\epsilon^{\perp}}$) were calculated over range of $\epsilon^{\perp} \in [-200, 200]$ with a grid size of 1, except in the range $\epsilon^{\perp} \in [-10, 10]$ where a grid size of 0.1 was used for better resolution. In all cases a significance threshold of $p < 0.05$ was used.

### 6 GSEA with Compositional Weighting

This section consists of three parts. In the first part it is proven that the LFC of an entity $d$ (i.e., $\theta_d$) used in GSEA-LFC is related to the GSEA-CLR input $\vartheta_d$ by

$$\vartheta_d = \theta_d - \text{mean}\,(\theta)\,.$$

In the second part we define GSEA with Compositional Weighting (GSEA-CW). We demonstrate that GSEA-CW is a generalization of GSEA-CLR and that GSEA-CW target estimands are scale invariant as they may be expressed as a function of log-contrasts. In the final part we discuss another possible GSEA-CW target estimand and explore the scientific question it implies.

#### 6.1 GSEA-CLR

Here it is proven that $\vartheta_d$ is related to $\theta_d$ through the equality $\vartheta_d = \theta_d - \text{mean}\,(\theta)$. We start by noting that $\vartheta_d$ is, by definition, equivalent to the LFC estimated using the Centered Log Ratio (CLR) assumption as defined in equation (4) in the main text, denoted here as $\theta_d^{clr}$. Expanding on equation (4) in the main text:

$$\begin{aligned}
\theta_d^{clr} &= \underset{n:x_n=1}{\text{mean}}(\log W_{dn}^{clr}) - \underset{n:x_n=0}{\text{mean}}(\log W_{dn}^{clr}) \\
&= \underset{n:x_n=1}{\text{mean}}\left[\log \frac{W_{dn}^{\parallel}}{G(W_{\cdot n}^{\parallel})}\right] - \underset{n:x_n=0}{\text{mean}}\left[\log \frac{W_{dn}^{\parallel}}{G(W_{\cdot n}^{\parallel})}\right] \\
&= \underset{n:x_n=1}{\text{mean}}\left[\log W_{dn}^{\parallel} - \log G(W_{\cdot n}^{\parallel})\right] - \underset{n:x_n=0}{\text{mean}}\left[\log W_{dn}^{\parallel} - \log G(W_{\cdot n}^{\parallel})\right] \\
&= \underset{n:x_n=1}{\text{mean}}\left[\log W_{dn}^{\parallel}\right] - \underset{n:x_{n=1}}{\text{mean}}\left[\log G(W_{\cdot n}^{\parallel})\right] - \underset{n:x_n=0}{\text{mean}}\left[\log W_{dn}^{\parallel}\right] + \underset{n:x_{n=0}}{\text{mean}}\left[\log G(W_{\cdot n}^{\parallel})\right] \\
&= \left(\underset{n:x_n=1}{\text{mean}}\left[\log W_{dn}^{\parallel}\right] - \underset{n:x_n=0}{\text{mean}}\left[\log W_{dn}^{\parallel}\right]\right) + \underset{n:x_{n=0}}{\text{mean}}\left[\log G(W_{\cdot n}^{\parallel})\right] - \underset{n:x_{n=1}}{\text{mean}}\left[\log G(W_{\cdot n}^{\parallel})\right] \\
&= \theta_d^{\parallel} + \underset{n:x_n=0}{\text{mean}}\left[\log G(W_{\cdot n}^{\parallel})\right] - \underset{n:x_n=1}{\text{mean}}\left[\log G(W_{\cdot n}^{\parallel})\right] \\
&= \theta_d^{\parallel} + \underset{n:x_n=0}{\text{mean}}\left[\underset{j=1,\ldots,D}{\text{mean}}\left(\log W_{jn}^{\parallel}\right)\right] - \underset{n:x_n=1}{\text{mean}}\left[\underset{j=1,\ldots,D}{\text{mean}}\left(\log W_{jn}^{\parallel}\right)\right] \\
&= \theta_d^{\parallel} + \underset{j=1,\ldots,D}{\text{mean}}\left[\underset{n:x_n=0}{\text{mean}}\left(\log W_{jn}^{\parallel}\right)\right] - \underset{j=1,\ldots,D}{\text{mean}}\left[\underset{n:x_n=1}{\text{mean}}\left(\log W_{jn}^{\parallel}\right)\right] \\
&= \theta_d^{\parallel} + \underset{j=1,\ldots,D}{\text{mean}}\left[\underset{n:x_n=0}{\text{mean}}\left(\log W_{jn}^{\parallel}\right) - \underset{n:x_n=1}{\text{mean}}\left(\log W_{jn}^{\parallel}\right)\right] \\
&= \theta_d^{\parallel} + \text{mean}\left(-\theta^{\parallel}\right) \\
&= \theta_d^{\parallel} - \text{mean}\left(\theta^{\parallel}\right).
\end{aligned}$$

To complete the proof it is shown that $\vartheta_d = \theta_d - \text{mean}(\theta)$ is equivalent to $\theta_d^{clr} = \theta_d^{\parallel} - \text{mean}\left(\theta^{\parallel}\right)$ because of the equality $\theta_d - \text{mean}(\theta) = \theta_d^{\parallel} - \text{mean}\left(\theta^{\parallel}\right)$:

$$\begin{aligned} \theta_d - \text{mean}(\theta) &= \left(\theta_d^{\parallel} + \theta^{\perp}\right) - \text{mean}\left(\theta^{\parallel} + \theta^{\perp}\right) \\ &= \theta_d^{\parallel} + \theta^{\perp} - \text{mean}(\theta^{\parallel}) - \theta^{\perp} \\ &= \theta_d^{\parallel} - \text{mean}(\theta^{\parallel}). \end{aligned}$$

Thus $\vartheta_d = \theta_d - \text{mean}(\theta) = \theta_d^{clr}$. Notably, because the scale term $\theta^{\perp}$ cancels out, $\vartheta_d$ may be calculated as $\theta_d - \text{mean}(\theta)$ from any estimated $\hat{\theta}$, regardless of the assumed value $\hat{\theta}^{\perp}$.

#### 6.2 GSEA-CW as a Function of a Log-contrast of $\theta$

Following our notation in the main text, let $\theta$ denote a vector of LFCs, $\theta^{\parallel}$ denote the corresponding vector of log-fold-changes in proportions, and $\theta^{\perp}$ be a scalar denoting the log change in the scale of the system between two experimental conditions. We can then write $\theta = \theta^{\parallel} + \mathbf{1}\theta^{\perp}$ where $\theta^{\parallel}$ is a vector of $D$ elements as defined in the main text and $\mathbf{1}$ is a vector with every element equal to 1.

A log-contrast of $\theta$ is a multivariate function $l : \Theta \to \mathbb{R}^P$ defined as

$$l(\theta) = \Psi\theta$$

for a $P \times D$ contrast matrix $\Psi$ satisfying the constraint

$$\sum_d \Psi_{pd} = 0 \quad \forall p \in \{1, \dots, P\}.$$

Note that log-contrasts are scale-invariant functions which we illustrate by showing that $l(\theta) = l(\theta^{\parallel})$:

$$\begin{aligned} l(\theta) &= \Psi\theta \\ &= \Psi\theta^{\parallel} + \Psi\mathbf{1}\theta^{\perp} \\ &= \Psi\theta^{\parallel} \\ &= l(\theta^{\parallel}) \end{aligned}$$

where we have used the fact that $\Psi\mathbf{1}\theta^{\perp} = 0$ which follows from the sum-to-zero constraint on $\Psi$. It follows that any real-valued function $f$ of a log-contrast, i.e., $f(l(\theta))$ is also scale invariant as $f(l(\theta)) = f(l(\theta^{\parallel}))$.

We define the GSEA-CW(-S) (where the -$S$ suffix implies sample-label rather than entity-label permutations) target estimand as GSEA using a ranking statistic of the form $R = f(l(\theta))$. Using the notation in the main text, the target estimand for GSEA-CW(-S) can be expressed as $\varphi_S = u(f(l(\theta)))$. To ensure scale invariance, we require that the method chosen to estimate LFCs ($\theta$) does not rely on the scale for estimation of $\theta^{\parallel}$. This requirement is included to avoid trivial scale reliance being introduced into GSEA-CW(-S). Together this is sufficient to prove that the GSEA-CW(-S) target estimand is scale invariant.

The GSEA-CLR(-S) target estimand has the required form for GSEA-CW(-S), which can be proven by constructing the matrix $\Psi$ as

$$\Psi_{pd} = \begin{cases} \frac{(N-1)}{N}, & \text{if } p = d \\ -\frac{1}{N}, & \text{otherwise.} \end{cases} \tag{4}$$

The $D$ length vector of GSEA-CLR(-S) inputs may then be expressed as $\vartheta = \Psi\theta$ and the GSEA-CLR(-S) target estimand is therefore $\varphi_S = u(\vartheta)$.

##### 6.3 Another GSEA-CW Target Estimand

There exists an infinitely larger family of GSEA-CW target estimands that each implies a different scientific question of interest. Here we explore one such target estimand: the *absolute value* of the mean distance of each entity to all other entities. This target estimand may be expressed as $\varphi_S = u(\lambda)$ where (for entity $d$) we define $\lambda_d$ as

$$\lambda_d = \left| \theta_d - \frac{1}{D} \sum_{i=1}^{D} \theta_i \right|.$$

The log-contrast for $\lambda$ can be expressed using the same matrix $\Psi$ as GSEA-CLR but with an absolute sign added: $\lambda = |\Psi\theta|$.

The scientific question implied by this target estimand differs from GSEA-CLR in one critical way. Consider an entity set for which half of the entities are highly depleted and half the entities are highly enriched relative to the mean LFC. For GSEA-CLR, half these entities would cluster at the beginning of the ranked list and half would cluster at the end. For this target estimand, however, the absolute value allows all the entities in the set to cluster together at the beginning of the ranked list. In this example, GSEA-CW(-S) using this ranking statistic would likely result in a lower p-value and higher ES compared to GSEA-CLR(-S). This target estimand therefore implies that we are interested not only in entity sets that are exclusively enriched or depleted relative to the mean, but also in entity sets where a portion of the entities are enriched and the other portion are depleted relative to the mean.

#### 7 Balances

In Supplementary Section 6 we defined a class of scale invariant target estimands based on GSEA [4] using log-contrasts. However, alternative scale invariant methods for DSA not based on GSEA have been recently proposed. Nguyen et al. [16] outline a scale invariant method for DSA based on the *balance* between entites in some set of interest and not in this set of interest. For geometric mean function $\mathrm{G}(\cdot)$, entity set indices

$S$, entities $d$, and study participant $n$, a balance is calculated as

$$f_S(W_{\cdot n}) \propto \log \frac{\mathrm{G}(W_{(d \in S)n})}{\mathrm{G}(W_{(d \notin S)n})}. \tag{5}$$

Their DSA method then evaluates a null hypothesis using entity-label permutations to test if $f_S(W_{\cdot n})$ is an unusually large or negative value within each participant $n$ for each set $S$.

At face value, this balance formulation is appealing: it is scale invariant and intuitive, equating differential enrichment or depletion to changes in the average abundance between set and non-set entities. However, as we demonstrate below it can be highly sensitive to outliers as might occur when an entity is incorrectly annotated and thus erroneously included in a set to which it does not belong.

#### 7.1 Sensitivity of Balances and GSEA-CLR to Outliers

Here we illustrate the sensitivity of balances by considering measurements of 11 entities from two participants. We define a matrix of abundances $W$ as

$$W = \begin{bmatrix} 308 & 305 & 301 & 119 & 107 & 107 & 103 & 100 & 96 & 92 & 4 \\ 5 & 5 & 5 & 5 & 5 & 5 & 5 & 5 & 5 & 5 & 5 \end{bmatrix}^T.$$

Let each of the two columns of $W$ (notice the transpose) represent individual participants $n = [1, 2]$. Let the indices of the entities in the set of interest be given by $S = \{1, 2, 3, 11\}$, meaning that the counts associated with $S$ when $n = 1$ are $(308, 305, 301, 4)$. When $n = 1$ the first three entities in $S$ have substantially enriched counts compared to the last entity which represents an outlier (potentially generated by an annotation error) with a count of only 4. However when $n = 2$ the entities in and outside of the set $S$ all have counts of 5. Yet the balances for both $n = 1$ and $n = 2$ are approximately zero, implying that $S$ is not enriched in both $n$. Balances thus do not capture what we argue is a clear distinction between the two participants in the context of $S$. GSEA-CLR is however able to capture the distinction between participants 1 and 2. Calculating the LFC as $\theta_d = \log(W_{d1}/W_{d2})$, the GSEA-CLR enrichment score is 0.98 which is close to the maximum score of 1, indicating a high degree of enrichment.

If the outlier is removed from the set (i.e., letting $S = \{1, 2, 3\}$) then GSEA-CLR remains fairly unchanged: the enrichment score increases only slightly to 1. In contrast, when the outlier is removed from $S$ the results implied by balances change substantially: when $n = 1$ the balance increases to 1.5 while the balance when $n = 2$ remains at 0. With this removal of the outlier, the balances now distinguish between the participants.

### References


1. Gatti, D. M., Barry, W. T., Nobel, A. B., Rusyn, I. & Wright, F. A. Heading down the wrong pathway: on the influence of correlation within gene sets. *BMC Genomics* **11,** 574–574 (2010).
2. Wu, D. & Smyth, G. K. Camera: A competitive gene set test accounting for inter-gene correlation. *Nucleic Acids Res.* **40,** 1–12 (2012).
3. Tamayo, P., Steinhardt, G., Liberzon, A. & Mesirov, J. P. The limitations of simple gene set enrichment analysis assuming gene independence. *Stat. Methods Med. Res.* **25,** 472–487 (2016).
4. Subramanian, A. *et al.* Gene set enrichment analysis: a knowledge-based approach for interpreting genome-wide expression profiles. *Proc. Natl. Acad. Sci. U.S.A.* **102,** 15545–15550 (2005).
5. Hung, J.-H. *et al.* Identification of functional modules that correlate with phenotypic difference: the influence of network topology. *Genome Biol.* **11,** R23 (2010).
6. Aran, D. *et al.* Comprehensive analysis of normal adjacent to tumor transcriptomes. *Nat. Commun.* **8,** 1–13 (2017).
7. Nixon, M. P., Letourneau, J., David, L., Mukherjee, S. & Silverman, J. D. Scale Reliant Inference. Preprint at http://arxiv.org/abs/2201.03616 (2022).
8. Li, W.-X. *et al.* Comprehensive tissue-specific gene set enrichment analysis and transcription factor analysis of breast cancer by integrating 14 gene expression datasets. *Oncotarget* **8,** 6775–6786 (2017).
9. Lonsdale, J. *et al.* The Genotype-Tissue Expression (GTEx) project. *Nat. Genet.* **45,** 580–585 (2013).
10. Rahman, M. *et al.* Alternative preprocessing of RNA-Sequencing data in the Cancer Genome Atlas leads to improved analysis results. *Bioinformatics* **31,** 3666–3672 (2015).
11. Liberzon, A. *et al.* Molecular signatures database (MSigDB) 3.0. *Bioinformatics* **27,** 1739–1740 (2011).
12. Ritchie, M. E. *et al.* limma powers differential expression analyses for RNA-sequencing and microarray studies. *Nucleic Acids Res.* **43,** e47–e47 (2015).
13. Thorpe, L. E. *et al.* Rationale, design and respondent characteristics of the 2013–2014 New York City Health and Nutrition Examination Survey (NYC HANES 2013–2014). *Prev. Med. Rep.* **2,** 580–585 (2015).
14. Beghini, F. *et al.* Tobacco exposure associated with oral microbiota oxygen utilization in the New York City Health and Nutrition Examination Study. *Ann. Epidemiol.* **34,** 18–25.e3 (2019).
15. Berger, R. L. & Boos, D. D. P values maximized over a confidence set for the nuisance parameter. *J. Am. Stat. Assoc.* **89,** 1012–1016 (1994).
16. Nguyen, Q. P., Hoen, A. G. & Frost, H. R. CBEA: Competitive balances for taxonomic enrichment analysis. *PLoS Comput. Biol.* **18,** 1–24 (2022).
